# Rational decision self-organizes beyond subjective reasonings through the rhythmic coupling of brain and internal organs

**DOI:** 10.1101/2025.03.05.641768

**Authors:** Shogo Yonekura, Hoshinori Kanazawa, Takuto Tsuji, Atsushi Narita, Yasuo Kuniyoshi

## Abstract

Psychological evidence shows that updating the rhythmic, internal body activity (IBA) based on the decision outcome is important to have a rational decision-making in a gambling task. However, the detailed mechanism how body guides rationality remains largely unclear. This paper provides a comprehensive theory that bridges the rhythmic coupling between the brain and internal organs and the rationality in a gambling decision-making task. Through our study of human binocular rivalry, we show that IBA creates the decision-switching (Sw) and -locking (Loc) phases based on the IBA parameters such as frequency, amplitude, and mean activity. When applied to a gambling task, the Sw-Loc structure creates directionality within the expected-value space of the decision options, wherein only options with expected values aligned with that directionality are stabilized. The stable direction can flip by changing how the IBA is updated in response to a decision outcome (e.g., taking a deeper or shallower breath). Using a theoretical analysis of a Brownian particle in a two-state heat bath, we show that the Sw region induces a large first-passage decision bias based on the random decision switching rate. Then, the Loc region acts as a rectifier, allowing only options with expected values aligned with the stable direction to persist. The rhythmic coupling of a brain and internal organs creates a directed medium that absorbs and separates external information, inducing biased decision-making through randomness.

## 1 Introduction

The somatic marker hypothesis (SMH) suggests that information from an animal’s internal organs is necessary for the brain to make rational decisions [14, 16, 15, 5]. A meta-analysis of studies on SMH strongly supports the idea that bodily states in response to decision outcomes guide overall performance in gambling tasks [51]. However, the idea that the body can anticipate and represent the value of each decision option before the brain (or conscious awareness) is only weakly supported (or rejected by several studies) [51, 42, 18, 61]. A recent study points out that the effect of the bodily response may reverse from warning about risky options to favoring safe ones with continued trials. Therefore, this dynamic characteristic of somatic markers may mask or reduce statistical significance [20]. In any case, contrary to SMH’s original claim, it is reasonable to assume that the body does not merely predict or represent the expected value of each decisional option before the brain does. Nevertheless, the body still influences the decision-making process. Therefore, the precise mechanism by which the body influences rational decision-making remains unclear.

On the other hand, recent research shows that internal bodily activities (IBAs) can influence brain and cognitive activity by rhythmically coupling with the brain via interoception or neuro-modulation, and this rhythmic coupling affects cognitive processes such as memory and perception [25, 24, 63, 55, 49, 59, 46, 28, 37, 23, 1, 21, 57, 19]. These studies prompt us to reconsider SMH in terms of the relationship between rhythmic IBAs and rational decision-making processes. If we can explain SMH – that rationality is guided by the body – without assuming that a priori, meta-physical reward and value information is embedded in our internal organs and the brain merely receives value representations from them [16], then that model must be physically fundamental. Understanding this physically fundamental decision-making mechanism will provide a more sophisticated understanding of how metaphysical information such as reward and value exist in living creatures.

Our initiative consists of the following four steps.

1. Firstly, we study how rhythmic IBAs influence decision-making based on human binocular rivalry paradigm. We studied binocular rivalry because the underlying neural circuit is isomorphic to the general (behavioral) decision-making [32, 38, 56, 2, 9, 7]. Therefore, studying binocular rivalry enable us to examine the physical effect of IBA while abstracting away metaphysical information such as reward and value. We demonstrate that human cardiorespiratory activity induces spontaneous switching of perceptual decisions.
2. Secondly, we conduct computational and dynamical analysis for the core dynamics of how IBAs influence neural circuit involved in decision-making. We demonstrate that rhythmic IBAs induce bifurcations of the decision neural circuit into decision-locking (Loc) and -switching (Sw) modes based on IBA parameters.
3. Thirdly, we use a computational simulation of the Iowa Gambling Task (IGT) [5], which was designed by Bechara and Damasio, to test human rationality in a behavioral decision-making task. In IGT, there are two types of options: one with low score variance and positive expected value (EV) and one with high variance and negative EV. In IGT, whether participants can choose an profitable deck is considered the criterion for rationality. We demonstrate that a neural decision-making circuit can flip the bias toward either profitable or unprofitable gambling options based on how it updates the IBA according to the outcome of the decision.
4. Fourth, we propose a theory in which the decision-making process in the Sw region is viewed as a particle transport process. In this model, decision options compete to control a particle in IBA parameter space in order to reach the Loc region. In particular, we propose a model of a particle driven by a two-state heat bath that switches randomly, where the heat bath states correspond to stochastic dynamics with different non-zero drift terms and variances of decisional options, and a particle moves between the homeostatic lower reflecting boundary and the upper absorbing boundary. We derive the exact expression of decision-bias using the survival function [11, 6, 40], and present how decision bias is induced and flipped in the Sw region.

Understanding the bias-generating mechanism in the Sw region enables an understanding of the multilayered mechanisms underlying the biases formation across the entire Sw-Loc region: The rhythmic coupling of the brain and IBA creates a value space where the direction of value is determined by how the IBA updates in response to the outcome of a decision. The Sw region then induces the first-passage decision bias, while the Loc region permits only valuable options.

Throughout this paper, we demonstrate how internal organ functions operate at different levels to induce decision bias and coordinate and align subjective value functions with those of others to achieve intersubjective rationality.

## 2 Human binocular rivalry is influenced by internal organs

In a human binocular rivalry experiment, we measured respiratory chest motion, electrocardiogram (ECG), and pupil diameter under natural breathing conditions. The phase of chest movement *θ*(*t*) corresponding to the respiratory phase, the cardiac R-wave *R*(*t*), the normalized pupil diameter *d*_*p*_(*t*), the blink timing *B*_*l*_(*t*), the perceptual switching time *e*_*p*_ were calculated (Fig. 1-(a, b), Appendix A).

**Figure 1:**
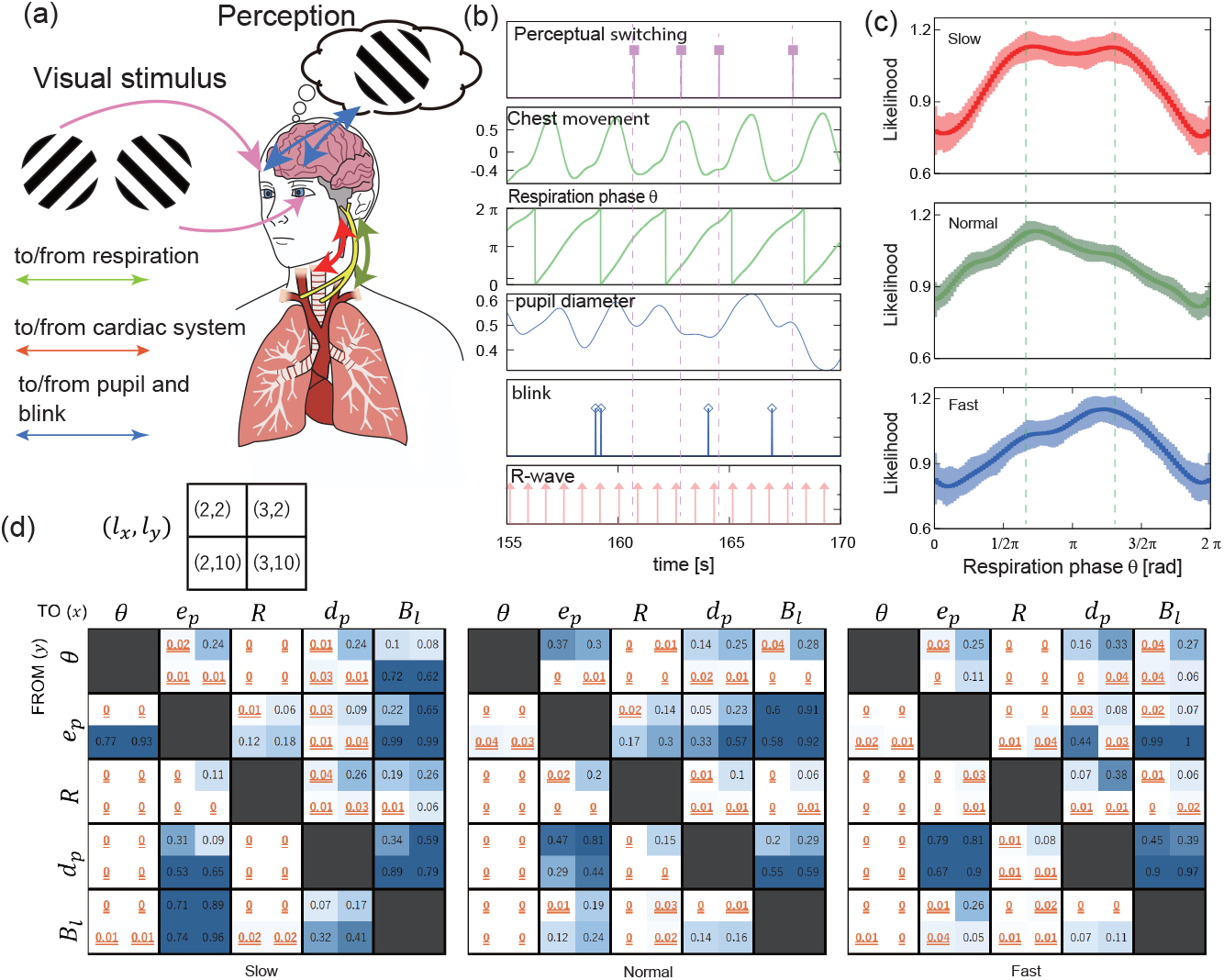
Internal organs influence perceptual alternation. **(a)** A schematic of human experiment based on the binocular rivalry paradigm. We measured respiratory chest movement, electrocardiogram, and pupil diameters, and the timing of perceptual alternation. **(b)** A sample of the perceptual alternation timing *e*_*p*_(*t*), the phase of chest movement *θ*(*t*), the cardiac R-wave *R*(*t*), the normalized pupil diameter *d*_*p*_(*t*), and the blink timing *B*_*l*_(*t*). **(c)** The estimated likelihood of perceptual alternation in human participants with respect to the respiration rate (slow, normal, fast) and the respiration phase *θ*, 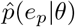 with the 95 % highest density interval (HDI). (**d**)Heatmaps showing the information flow from/to *θ, e*_*p*_, *R, d*_*p*_, and *B*_*l*_. The 2 × 2 submatrix denotes the embedding dimension *l*_*x*_, *l*_*y*_ used to compute TE from *y*→ *x*. The p-values which satisfy the condition *<* 0.05 are indicated by red text with a double underline.x

The likelihood of a perceptual alternation event, *e*_*p*_, given the respiratory phase *θ* and rate (slow, normal, fast) was estimated using the Markov Chain Monte Carlo (MCMC) method [12] on data from 30 male participants (mean age is 25.5 with a standard deviation of 2.8). Fig. 1-(c) shows the mean and 95 % highest density interval (HDI) of the estimated likelihood function 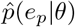. It is evident that 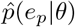 depends on the respiration phase *θ*, and that the shape of 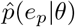 is influenced by the respiration rate.

To confirm the statistical significance of the causal effects between IBAs and perceptual alter-nation, we analyzed information flow using transfer entropy (TE). In particular, TE between *e*_*p*_, *θ, R, d*_*p*_ and *B*_*l*_ are estimated using a method for calculating TE between point processes [50]. The variables *θ*(∈ [0, 2*π*]) and *d*_*p*_(∈ [0, 1]) of the nonpoint process have been converted to point processes (see Appendix-A). The heatmap in Fig. 1-(d) shows the p-values obtained from the permutation tests using the point-process TE described in Ref. [50]. P-values less than 0.05 are indicated by red, double-underlined text. The results indicate that there is a significant flow of information from the respiratory phase and the cardiac R-wave to the perceptual alternation.

The vector of embedding dimensions (*l*_*x*_, *l*_*y*_) is a parameter that indicates how many spikes are used to compute the TE from the source process *y* to the target process *x*. The respiratory phase *θ* has a significant information flow with *l*_*y*_ ≥ 10 with *x* = 2, 3 in slow and normal breathing rates. (The point process *Y* corresponding to the respiration phase *θ* with *I*_*y*_ = 10 denotes one cycle of respiration; for we computed the point process by dividing the parameter space *θ* : [0, 2*π*] into 10 subspaces. See also Appendix-A). This implies that one cycle of breathing has a significant contribution to two and three perceptual switching events, in slow and normal breathing rates. In contrast, at fast breathing rate, TE was only significant for *I*_*x*_ = 1, 2. This implies that at fast breathing rate, breathing has only a short term influence. (See Appendix-A and Fig. 10–12 for the explanations of parameters and discussions of parameter sensitivity). Overall, these results demonstrate that IBAs and perceptual alternation exert a significant influence on one another. The following demonstrates that spontaneous decision switching is an essential characteristic of the coupled brain-IBA system.

## 3 Internal organs can force the decisions to randomly-switch or lock

It has been demonstrated that respiration entrains the overall activity of cortical neurons [63, 37, 1] based on the following information pathways; (a) neuromodulation of cortical neurons by the locus coeruleus, which receives respiratory information (*g*(*t*)) [43, 47, 41, 48, 17, 45, 62], and (b) the broadcast of respiratory sensory information to cortical neurons via the olfactory bulb [63, 27, 34] (*A*(*t*)) (Fig. 2-a). However, it is still unclear how rhythmic entrainment induces decision alternations. The minimal model that can account for the perceptual alternation observed in binocular rivalry is a stochastic bistable system [35, 13, 9, 22]. To understand the core mechanism by which IBAs influence decision-making, we therefore conducted numerical simulations and mathematical analyses using the following minimal stochastic bistable neural model (see also Fig. 2-(b), Appendix-B):

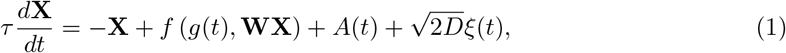

where matrix **X** denotes the membrane state of neurons, matrix **W** represents the synaptic connections, function *f* (*g*(*t*), *x*) = 1*/* (1 + exp (−*g* (*t*) *x*)) represents the activation function of neurons, and 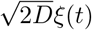 denotes the Gaussian noise with the standard deviation of *D*. This winner-take-all (WTA) motif circuit is seen not only for perceptual alternation but also for behavioral level [32, 38, 56, 2, 9]. The effect of IBAs via neuromodulation is given by *g*(*t*) = *g*_*g*_ sin 2*πft* + *g*_*b*_, and the effect via additive sensory input is given by *A*(*t*) = *A*_*g*_ sin 2*πft* + *A*_*b*_ (Fig. 2-b, Appendix-B). Fig. 2-(c) shows the switching of a winner neuron occurs in synchronization with the IBA-driven *g*(*t*), confirming that IBAs can directly induce decision switching.

**Figure 2:**
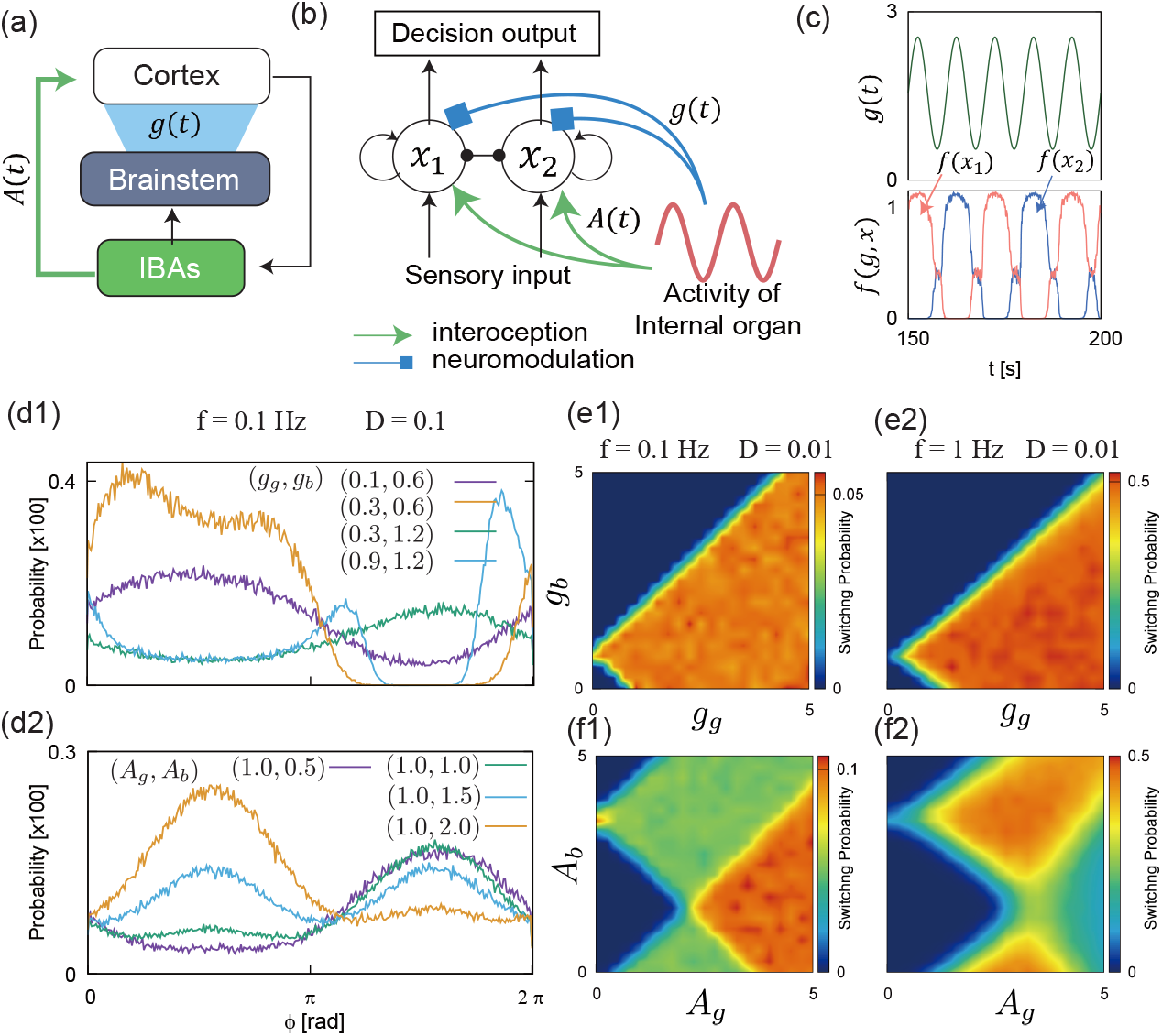
Internal organs create Loc/Sw modes in a neural decision-circuit. **(a)** Information from the animal IBA is possibly passed to cortical neurons via two pathways: neuromodulation *g*(*t*) and interoceptive sensory input *A*(*t*). **(b)** The simplest bistable neural model of perceptual or behavioural decision-making copled with IBA. IBA information is passed to competing neurons *x*_1_ and *x*_2_, via sensory input *A*(*t*) and neuromodulation *g*(*t*). Each competing neuron has a self-excitatory connection. **(c)** The winning neural outputs, *f* (*x*_1_) and *f* (*x*_2_), are randomly switched in a synchronised manner with periodic neuromodulation, *g*(*t*). **(d1, d2)** The switching probability of winning neurons *p*(*e*_*p*_| *ϕ*) under the influence of *g*(*t*) and *A*(*t*). The winning neuron tends to switch stochastically at a certain phase of *g*(*t*) and *A*(*t*), either at an angle of *π/*2 or 3*π/*2. **(e1, e2, f1, f2)** The heatmap of switching frequency of winning neuron [Hz] in relation to neuromodulation bias (*g*_*b*_) and gain (*g*_*g*_), and sensory bias (*A*_*b*_) and gain(*A*_*g*_).

Fig. 2-(d1, d2) demonstrates that the switching of the winning neuron is likely to occur at a specific phase *ϕ* of IBA. Furthermore, the *ϕ* that maximizes *p*(*e*_*p*_|*ϕ*) shifts with the IBA parameters *f, g*_*b*_, *g*_*g*_, *A*_*b*_, and *A*_*g*_.

The parameter space of IBA where decision-switching occur is linear with respect to *g*_*b*_ and *g*_*g*_ (neuromodulator bias and gain), and nonlinear with respect to *A*_*b*_ and *A*_*g*_ (sensory bias and gain) (Fig. 2-e1–f2).

### Spontaneous flip of the feedback modes is a core mechanism of decision switching

A simple mathematical analysis can demonstrate the core mechanism by which IBA influence decision-making. Taking the Taylor expansion of Eqs. 1 with *x*_*i*_ ∼ 0, and computing *x*_1_ −*x*_2_, we immediately obtain the one-dimensional linear differential equation with respect to 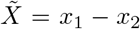 as

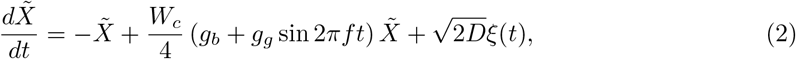

where *W*_*c*_ = *w*_*exc*_ − *w*_*inh*_, and *w*_*exc*_ and *w*_*inh*_ are the weights for the excitatory self-coupling and the mutual inhibitory connection, respectively (Appendix-B). It is shown that the dynamics of 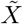 is inverted from negative feedback to positive feedback based on the phase of the periodic neuromodulator, this is consistent with the results in Fig. 2-(d1). It is mathematically shown that the switching to the positive feedback mode occurs when −1 + 1*/*4*W*_*c*_ (*g*_*b*_ + *g*_*g*_ sin 2*πft*) *>* 0, i.e., *g*_*b*_ *>* 4*/W*_*c*_ − *g*_*g*_ sin 2*πft*. The function *g*_*g*_ sin 2*πft* sweeps the range [−*g*_*g*_, *g*_*g*_], and a sufficiently large *g*_*b*_ blocks the transition of winning neurons. Therefore, the decision switching occurs around the range shown in Fig. 2-(e1,e2).

In contrast, the effect of interoceptive sensory input is highly nonlinear, as seen in Fig. 2-(f2). The nonlinear effect of *A*(*t*) to decision-making can be analyzed by using Itô’s formula [29]. If the activation gain *g*_*b*_ is sufficiently large, then the difference in neural outputs, *F* (*x*_1_, *x*_2_) = *f* (*x*_1_) − *f* (*x*_2_), is described as

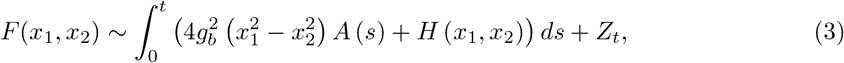

where *H*(*x*_1_, *x*_2_) is a function of *x*_1_ and *x*_2_, *Z*_*t*_ is a noise term. It is shown that *F* (*x*_1_, *x*_2_) receives the quadratic negative feedback effect 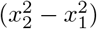 when *A*(*t*) is in the phase [0, *π*] and similarly, the positive feedback effect in the phase [*π*, 2*π*]. For the detailed derivation and explanation of Eq. 3, see Appendix-B.

## 4 Decision-bias self-organizes

In IGT, an agent repeatedly draws a card from a bad (BD) or good (GD) deck, where the expected value of each deck is unknown. BD is designed to generate a high random score in the short term. However, the expected return is negative, so selecting BDs results in long-term losses. In contrast, GD is configured to produce only a small random score, but it has a positive expected return.

Thus, choosing GDs leads to long-term profits. The score *R*_*n*_, that an agent receives at the n-th decision is designed as follows: 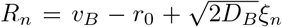for BD and 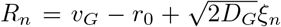 for GD, where *ξ*_*n*_ is a random number generated from a normal distribution. We used *v*_*B*_ = −25, *v*_*G*_ = 25, 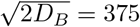, and 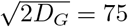 in Fig. 3.

**Figure 3:**
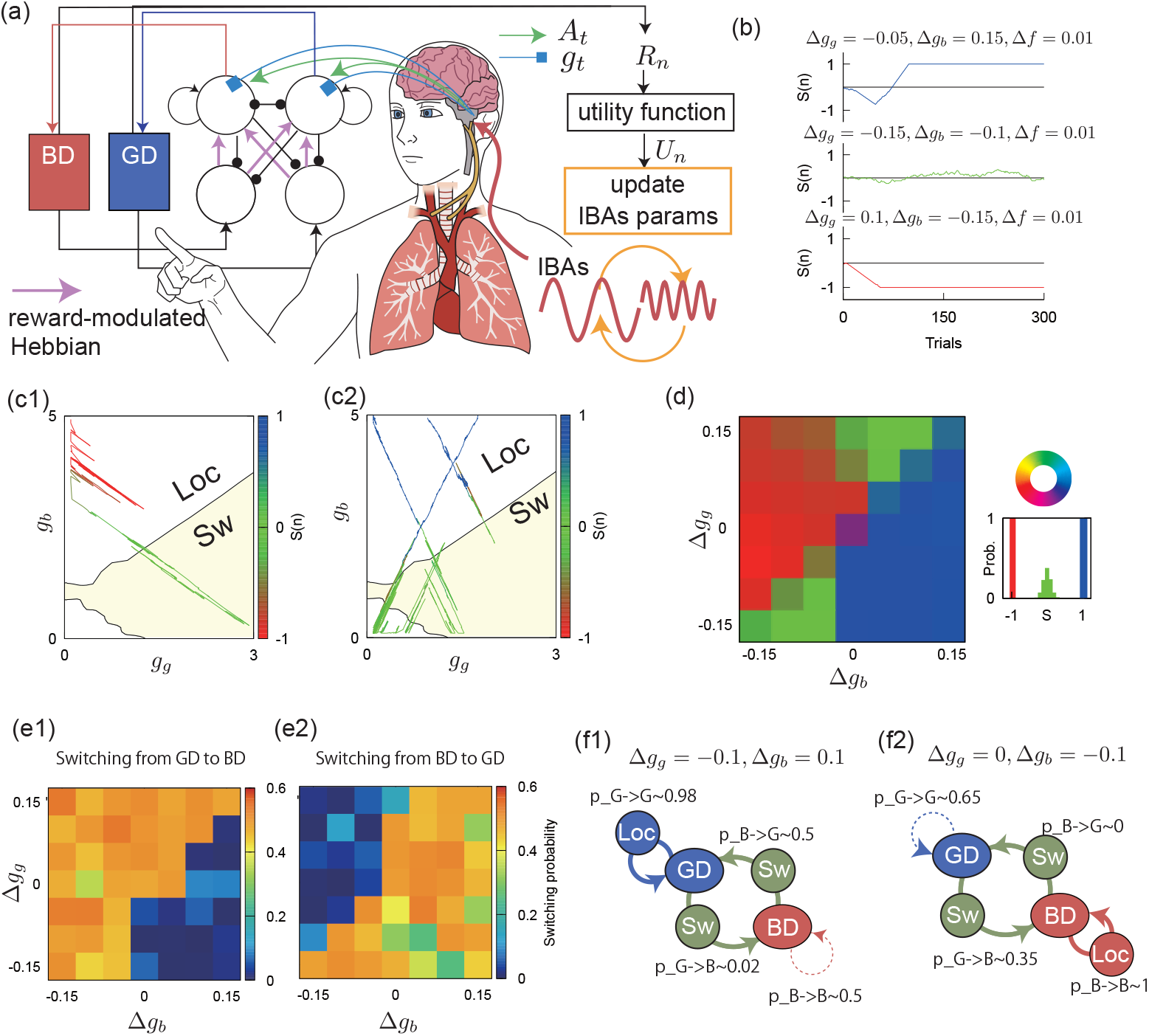
A minimal model of somatic marker in IGT. **(a)** A schematic diagram of the task and a computational somatic marker model. BD is bad deck and GD is good deck. The decisional neural circuit repeatedly chose one of the decks and received the score *R*_*n*_. *R*_*n*_ is evaluated internally, and IBAs are updated based on *U*_*n*_. The decision neural circuit is basically the same with Fig. 2 but has top-down inhibitory connection, which is called “evidence cut-off” [9]. The task performance is measured by *S*(*n*), which represents how often the agent selected GD. **(b)** Trajectories of *S*(*n*) when the agent successfully converged to GD *S*(∞) = 1, and BD *S*(∞) = −1, or keep wondering *S*(∞) ~ 0. **(c1, c2)** Trajectories of *g*(*t*) for *S*(∞) = −1 **(c1)** and *S*(∞) = 1 **(c2)** are superimposed on the phase diagram of Loc and Sw. **(d)** The influence of the mapping functions of *U*_*n*_ → Δ*g*_*g*_, Δ*g*_*b*_. We computed the histogram of *S*(∞), i.e., *p*(*S*(∞) = *x*) for each parameter set (Δ*g*_*b*_, Δ*g*_*g*_), and colored it according to the ratio of red for *p*(*S*(∞) = −1), green for *p*(*S*(∞) = 0), and blue for *p*(*S*(∞) = 1). **(e1, e2)** The transition probabilities from GD to BD and from BD to GD, respectively. **(f1, f2)** Understanding the entire system using finite state machines; the switching probability from GD →GD via Loc mode, GD →BD via Sw mode are measured using several typical parameters.

We use the measure *S*(*n*), which denotes how often an agent selects GD; If the agent consistently chooses GD, *S*(*n*) converges to 1; if the agent consistently chooses BD, *S*(*n*) converges to −1; and if the agent consistently switches between GD and BD, *S*(*n*) ∼ 0 (Fig. 3-a, Appendix-C).

We assumed that the neural gain is influenced by rhythmic IBAs as follows: *g* = *g*_*g*_ sin(2*πft*) + *g*_*b*_. The agent evaluates the outcome of the *n*th decision, *R*_*n*_, using the internal utility function *U*_*n*_ = *U* (*R*_*n*_); *U* (*R*) = *R* for *R >* 0 and −|*R*|^*β*^ for *R <* 0 (*β <* 1 means risk underestimation). After every decision, the IBA parameters are updated in a linear manner based on *U*_*n*_ as follows: *z*(*n* + 1) = (1 + *ζ*_*n*_)*z*(*n*) + Δ_*z*_(*λU*_*n*_ + *η*_*n*_), where *z* denotes either *g*_*g*_, *g*_*b*_, or *f, λ* is a scaling factor, and *η* and *ζ* are small Gaussian noise (Appendix-C).

Fig. 3-(b) show that the agent can reach a rational decision and converge through the IBA-induced spontaneous and implicit transitions between Loc and Sw (Fig. 3-c1, c2). Furthermore, the top panel in Fig. 3-(b) shows that the agent successfully performed spontaneous reversal learning, which is believed to be a key function of somatic markers in healthy persons; the agent first selected BD because it provided large rewards for the first ∼50 trials. However, BD soon started to return large negative scores, and the agent successfully changed its decision and converged on GD.

Fig. 3-(d) shows that metalearning, i.e., an appropriate configuration of Δ_*z*_ (i.e., Δ*g*_*g*_ or Δ*g*_*b*_) by a brain is necessary for the occurrence of rationality emergence. If Δ*z* is not adequate, the system is trapped to irrational behaviors (the bottom panel of Fig. 3-b, and c1).

The overall behavior of the coupled sytem is described as a finite state machine (Fig. 3-e, f); if *U*_*n*_ *<* 0 and *U*_*n*_ *>* 0 are mapped to IBA’s parameter update toward Loc and SW regions, respectively, then, the decision of BD is stabilized in this overall process because BD returns negative outcome in average. In contrast, if *U*_*n*_ *<* 0 and *U*_*n*_ *>* 0 are mapped to IBA’s parameter update toward SW and Loc modes, respectively, then, the decision is locked to GD in this overall process because GD returns positive outcome in average. It is important to note that the agent neither knows nor calculates the expected values of BD and GD.

Fig. 4 shows that for the asymmetric profit-loss stucture of IGT (*r*_0 ≠_ 0), the optimal configuration of Δ_*z*_ to spontaneously generate rational decisions varies. It should be noted that the reward-modulated Hebbian learning rule [39, 44, 36], which is an implementation of reinforcement learning based on Hebbian rule, is very weak to risk-underestimation, and the learning process easily corrups for *β <* 1. Despite the distorted utility function, SMH succeeds in making rational decisions, though a precise and flexible configuration of Δ_*z*_ seems to be required.

**Figure 4:**
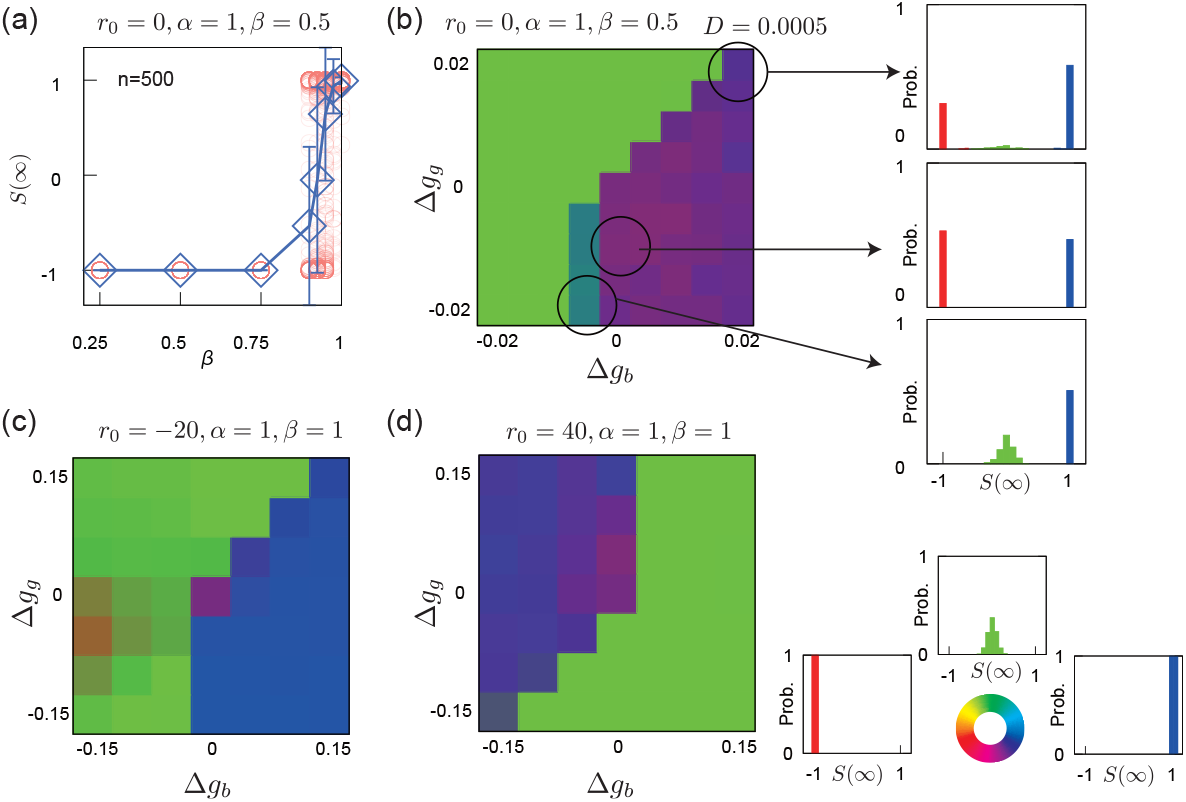
Reward-based Hebbian learing versus IBA-supported decision-making under distorted, asymmetric utility function. *U*_*n*_ **(a)** Limitation of reward-based Hebbian learning when the utiliy function underestimates risk (*β <* 1), the reinforcement learning easily drops to risky option; *S*(∞) = −1. **(b)** SMH can find rational decision *S*(∞) = 1 even with the highly risk underestimation *β* = 0.5, although such parameter region is narrow Δ − 0.005 ≤ *g*_*b*_ ≤ 0 and Δ*g*_*g*_ *<* 0. **(c)** For asymmetric bias *r*_0_ = −20 (the expected score of GD is large, i.e., = 45, whereas the expected score of BD is = −5), behaviors for *S*(∞) = −1 tends to disappear and most of behaviors converges to *S*(∞) = 0 or 1. **(d)** For *r*_0_ = 40, which means that both BD and GD return negative score in average, the parameter regions which yield the selection of GD are inverted compared to **(c)** and Fig. 3-(d).

## 5 First-passage decision bias

### 5.1 A particle in the two-state thermal bath

This section presents a theory explaining how decision biases can arise in the Sw region. We conceptualize the decision bias created in the Sw region as the first-passage bias and demonstrate that it originates from the interaction of random decision switching, a homeostatic lower-reflecting boundary, and an absorbing Loc boundary.

The trajectory of the IBA’s parameter (*g*_*g*_, *g*_*b*_) over decision trials shown in Fig. 3-(c1, c2), can be considered as a particle exhibiting the Brownian motion in the one-dimensional space *x*(*t*). Furthermore, the BD and GD decks can be considered as two different states of a thermal bath that impart different drift and diffusion forces to a particle. The stochastic dynamics are described as follows, with the notion that *z* describes the state of either GD or BD.

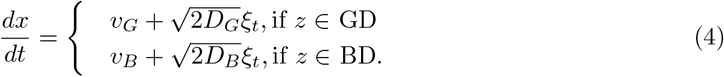

It is natural to introduce the lower reflecting boundary *x* = *L* and the upper absorbing boundary *x* = *H*: Sw regime is given by *x* : [*L, H*], and Loc regime is given by *x > H*. In the Sw regime, we can assume that the states of a thermal bath switch randomly at an instantaneous rate of *q*, and in the Loc regime, *q* = 0. In case of the numerical resutls shown in Fig. 3, we assumed that the neuromodulator *g*(*t*) *>* 0, and therefore *g*(*t*) = 0 corresponds to *L*, and the boundary of Sw-Loc corresponds to *T* .

To be precise, the BD → GD and GD → BD switching rates in the Sw region are not identical due to the noise terms *D*_*B*_ and *D*_*G*_. This should be checked by substituting *g*(*t, n*) = *g*_*b*_ + *g*_*a*_ sin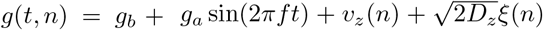 to the Eq. 1 and expanding it around *ξ* = 0 (*z* denotes either G or B, and *n* is the count of decision output). Nevertheless, we assume the the switching rates of BD →GD and GD →BD are identical for the simplicity. We demonstrate that first-passage state, that is, the thermal bath’s state when the particle first reaches the Loc (absorbing) boundary *x* = *H*, is largely biased even with identical decision switching rate *q*.

We can derive the coupled Fokker-Planck for this process as

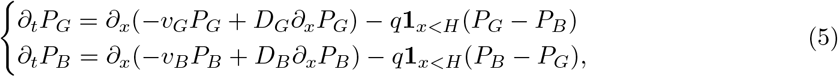

where *P*_*G,B*_(*x, t*) represents the probability that a particle is subject to the influence of GD or BD at position *x* at time *t*.

If *v*_*B*_ *<* 0, then, lim_*x→−∞*_ *P*_*B*_(*x, t*) = ∞ (if *L* → −∞). Likewise, if *v*_*G*_ *<* 0, then, lim_*x→∞*_ *P*_*G*_(*x, t*) = ∞ (if *H* → ∞). Thus, Eq. 5 can’t be solved directly.

The most general analytical approach for such divergent dynamics with *x* → ±∞ is to indroduce the survival probability *S*_*v*_(*t*) [11, 6, 40, 10]; *S*_*v*_(*t*) denotes how many particles are staying in the region *x* ∈ [*L, H*], as follows

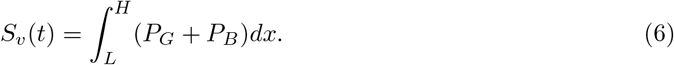

Furthermore, we assume that the particles which reach the absorbing boundary *x* = *H* are removed from the system. Then, *S*_*v*_(*t*) eventually become 0 as lim_*t→∞*_ *S*_*v*_(*t*) = 0. In virtue of *S*_*v*_(*t*), we can separate the space *x* and the time *t* as *P* (*x, t*) = *S*_*v*_(*t*)*µ*(*x*) and conduct the mode expansion of *P* (*x, t*) using the decay rate of *S*_*v*_(*t*). Here, we use the notation *P* (*x, t*) = (*P*_*G*_(*x, t*), *P*_*B*_(*x, t*))^*T*^ and similarly, *µ*(*x*) = (*µ*^(*G*)^(*x*), *µ*^(*B*)^(*x*))^*T*^ . Furthermore, when G and B appear in superscript or subscript, they represent the state of the heat bath (GD or BD).

### 5.2 Mode expansion using the decay rates *κ*_*n*_ of survival probability

In addition to the survival function *S*_*v*_(*t*), we introduce the variable *κ*(*t*)(∈ ℝ) which characterizes the decaying rate of *S*_*v*_(*t*) as 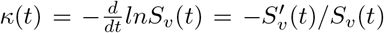. If *κ*(*t*) takes a constant value with *t* → ∞ using the linear combination of *κ*_*n*_ as

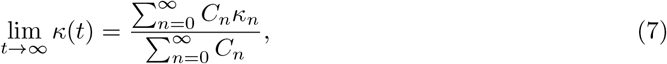

the survival probability *S*_*v*_(*t*) is represented as

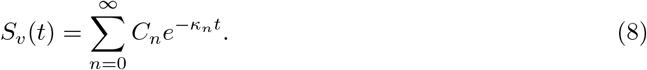

By using the representation of *S*_*v*_(*t*) in Eq. 8, we can separate time and position as

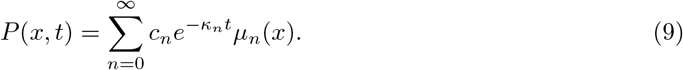

Note that we assumed *κ*_*n*_ is accompanied only by *µ*_*n*_. Furthermore, *C*_*n*_ in Eq. 8 is just a formal coefficient; *c*_*n*_ is the coefficient describing the contribution of each mode and is computed last in this section.

Each *µ*_*n*_ and *κ*_*n*_ satisfies the following Sturm-Liouville -type equation and boundary conditions

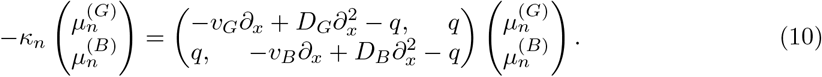

(Substituted Eq. 9 into the original FPE Eq. 5 yields this expression).

The reflecting and absorbing boundaries at *x* = *L, H* are described as

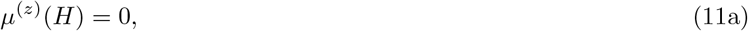

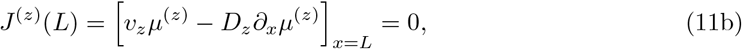

where *J* denotes flux.

All *κ*_*n*_ and *µ*_*n*_ must satisfy Eq. 10 and 11. If we further assume the *n*-th *µ*(*x*) is described as

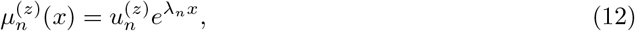

Eq. 10 is further simplified to the following eigenvalue problem

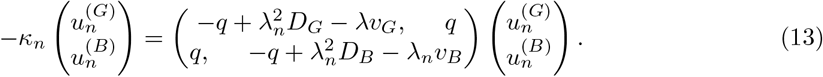

Now we describe Eq. 13 as 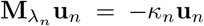. For the Eq. 13 has nontrivial solution, **u**_*n*_, 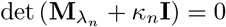 must be satisfied. Thus, *λ*_n_ and the ratio of eigenvectors 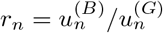 are computed by solving det 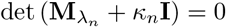. *r*_*n,i*_ is described as

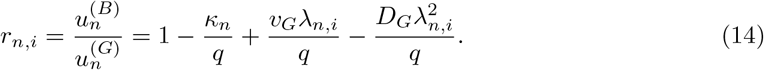

The analytical form of *λ*_*n*_ is too complex to list here, but is described as a function of *κ*_*n*_, *v*_*G,B*_,*D*_*G,B*_, *q, H*, and *L*. Now, the general solutions of 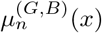are described as

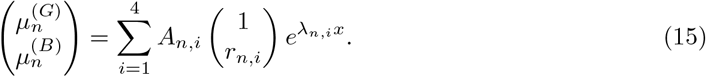

Computing *κ*_*n*_ and *A*_*n,i*_ requires boundary conditions given by Eq. 11. Substituting Eq. 15 yields the equation which *n*-th mode of *κ* and **A** must satisfy:

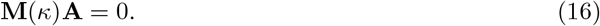

For the Eq. 16 has nontrivial **A**, det **M**(*κ*) must be zero. By solving det **M**(*κ*) = 0, we can compute discrete set of *κ* as *κ*_0_, *κ*_1_, · · ·, *κ*_*n*_, · · · . The coefficient *A*_*n*,1_ ∼ *A*_*n*,4_ which is correspondent to *κ*_*n*_ are computed as the null space of **M**(*κ*_*n*_). As far as we tested, there was no multiple root for *λ* whcih satisfied det **M**(*κ*) = 0.

To have the full description of Eq. 9, we need to compute the coefficient *c*_*n*_. This requires Kolmogorov backward equation (KBE) and consideration the biorthogonality of left and right eigen functions. The FPE in Eq. 10 is also described using the operator ℒ as ℒ *µ*_*n*_ = −*κ*_*n*_*µ*_*n*_. The adjoint equation is describe using the adjoint operator ℒ ^*†*^, and the eigenfunction *ψ*_*n*_ as ℒ ^*†*^*ψ*_*n*_ = −*κ*_*n*_*ψ*_*n*_ where

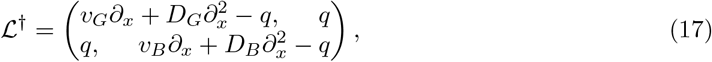

and *ψ*_*n*_(*x*) satisfies the following boundary equations *ψ*^*B*^(*H*) = 0, *ψ*^*G*^(*H*) = 0. *∂*_*x*_*ψ*^*B*^|_*x*=*L*_ = 0, and *∂*_*x*_*ψ*^*G*^|_*x*=*L*_ = 0. Intuitively, the right eigenfunction *µ*_*n*_(*x*) construct a probability density and the left eigenfunction *ψ*_*n*_(*x*) is the sensing function of the contribution of *µ*_*n*_(*x*). Eqs. 10 and 17 are Sturm-Liouville equations and share the same eigenvalues *κ*. Therefore, right and left eigenfunctions *µ*_*m*_ and *ψ*_*n*_ are biorthogonal as

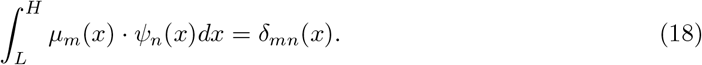

This biorthogonality enables the computation of *c*_*n*_ from the initial distribution *P* (*x, t* = 0) as

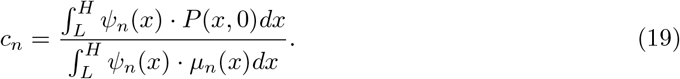

(This is because *P* (*x*, 0) =Σ _*n*_ *c*_*n*_*µ*_*n*_, and thus, 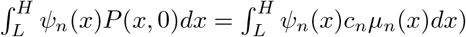.

In particular, the minimum *κ*_0_ and the correspondent eigenfunction 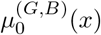 compose QSD. The higher order *κ*_*n*_ and *µ*_*n*_ describes the converging dynamics from the initial distribution *P* (*x*, 0) to *µ*_0_.

### 5.3 Measurements for the first-passage bias

In theoretical analysis, the decision bias can be characterized by using several measurement; absorbed fluxes *J*_*G*_(*x*_0_, *t*) and *J*_*B*_(*x*_0_, *t*), and the total escaping probability as GD, 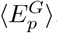 . Observing fluxes and their ratio are suitable to characterize the spatiotemporally-local decision biases which disappears by averaging over time and initial positions. On the other hand, 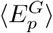is suitable to observe the global decision bias which never disappears even by averaging over time and initial positions.

The exact time evolution of the flux escaping from the absorbing boundary *x* = *H* is computed as

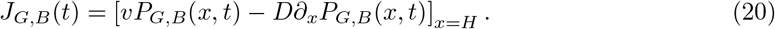

If we assume that the initial distribution is given by the Dirac’s delta function as *P*_*G*_(*x*, 0) = *P*_*B*_(*x*, 0) = 1*/*2*δ*(*x* − *x*_0_), the *c*_*n*_ becomes a function of *x*_0_ and we have

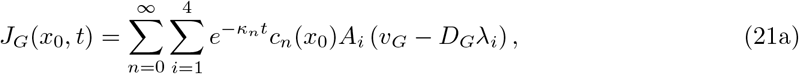

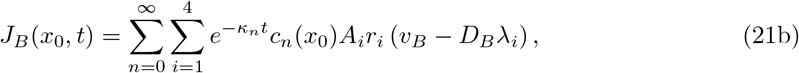

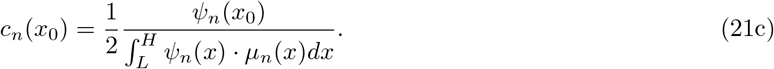

We have a similar expression of *J*(*x*_0_, *t*) when the initial distribution is given by the Gaussian distribution. The expression in Eq. 21 implicates that the resulting decision bias varies based on the initial position *x*_0_ and the time *t*. If the initial position *x*_0_ is sampled from the probability distribution function *P*_0_(*x*_0_), the average flux over the initial position *x*_0_, ⟨ *J*_*G,B*_(*t*) ⟩, is described as

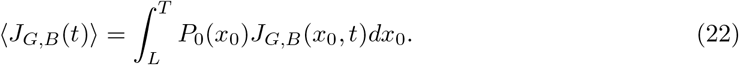

Fig.5-(a2, b2) shows theoretically-predicted and numerically-computed ⟨*J*_*G,B*_(*t*)⟩ obtained by sampling *x*_0_ from Gaussian distributions. It is shown that a temporally-partitioned assymmetry arises between ⟨*J*_*G*_(*t*)⟩ and ⟨*J*_*B*_(*t*)⟩. For example, in the panel (a2), *J*_*G*_ *> J*_*B*_ with *t* ∼ 0, but *J*_*G*_ *< J*_*B*_ with *t* → ∞. However, in several parameter regions, such asymmetry of *J*_*G*_ and *J*_*B*_ disappears by taking average of *J*_*G*_(*x*_0_, *t*) and *J*_*B*_(*x*_0_, *t*) over the time and the initial positions as

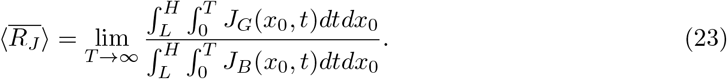

It should be noted that the assymetry of the fluxes shown in Fig. 5-(a2, b2) is almost recovered at the global level, as 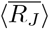takes a very close to 1; 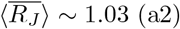 and∼1.005 (b2).

**Figure 5:**
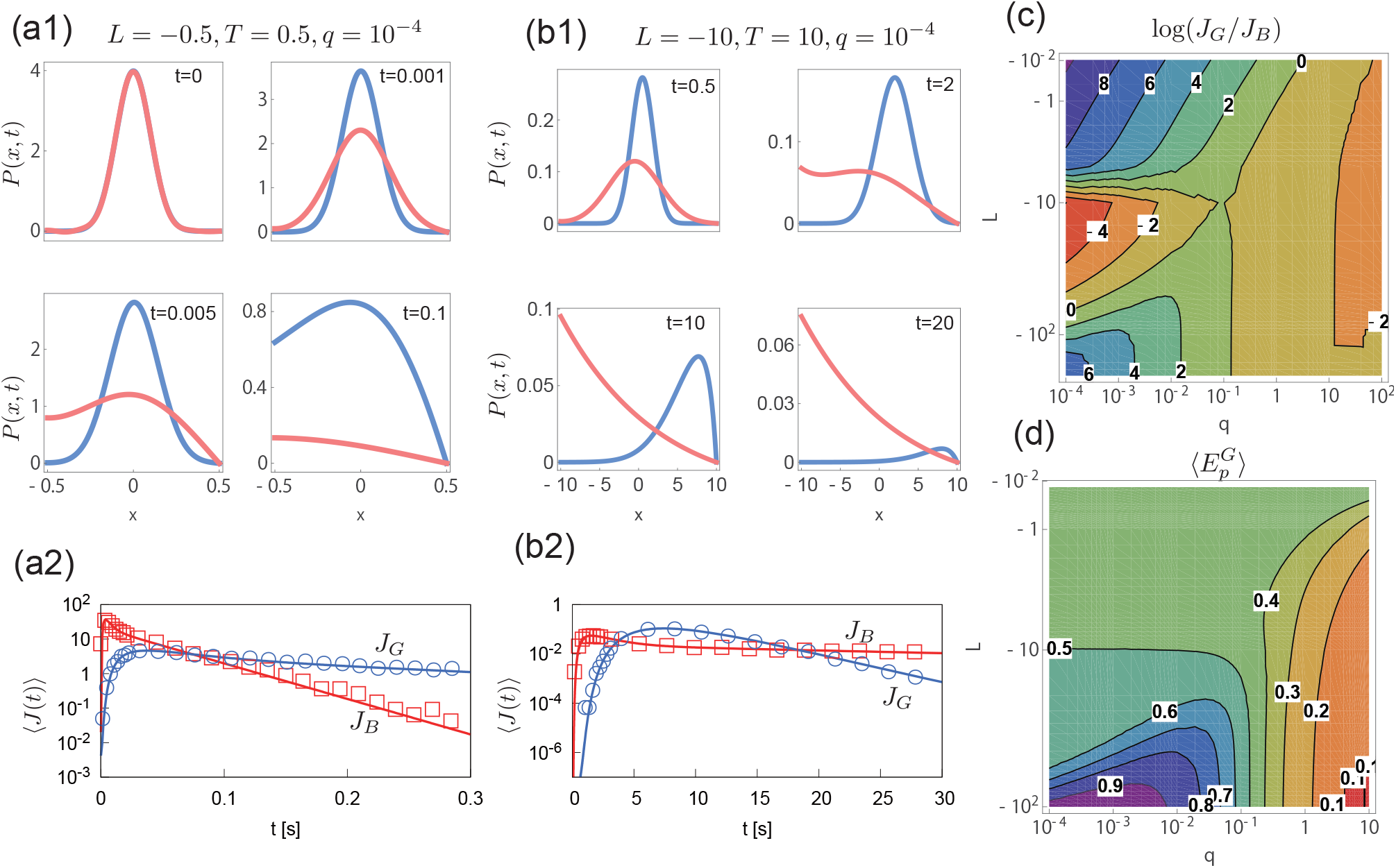
Temporally-partitioned and globally-persistent first-passage decision bias. **(a1, a2, b1, b2)** *P*_*G*_(*x, t*) (blue) and *P*_*B*_(*x, t*) (red), and *J*_*G*_(*x*_0_, *t*) (blue) and *J*_*B*_(*t, x*_0_) (red) with | *H* − *L* |= 1 and *q* = 10^*−*4^, for the panels (a), and | *H* − *L* |= 20 and *q* = 10^*−*4^ for the panels (b). Initial distribution is given by the Gaussian distribution with *x*_0_ = 0 with the standard deviation *σ* = 1*/*10 for (a), and *σ* = 1 for (b). In (a2), the bias exists temporally as ⟨*J*_*B*_⟩ ≫ ⟨*J*_*G*_⟩ for *t* ∼ 0 and ⟨*J*_*G*_⟩ ≫ ⟨*J*_*B*_⟩ for *t* ≫ 0. However, the total flux 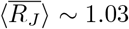, and the bias almost disappears at the global level. Similarly in (b2), *J*_*B*_ ≫ *J*_*G*_ for *t* ∼ 0, *J*_*G*_ ≫ *J*_*B*_ for *t* ∼ 10, and *J*_*B*_ ≫ *J*_*G*_ for *t* → ∞, but the total flux ratio 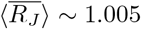 **(c)** The flux ratio log(*J*_*G*_*/J*_*B*_) in QSD, i.e., decision-bias in *t* → ∞. **(d)** The globally-persistent decision bias. It is clearly shown that the parameter region |*H* − *L*| *<* 10 and *q* ≪ 1 generates the locally-partitioned asymmetric decision bias and therefore 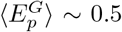 despite the *J*_*G*_*/J*_*B*_ in QSD is far from 1 (e.g., *log*(*J*_*G*_*/J*_*B*_) *>* 4 or *<* −4), and we need larger | *H* − *L* | or *q* ≫ 1 for the globally-persistent bias.

**Figure 6:**
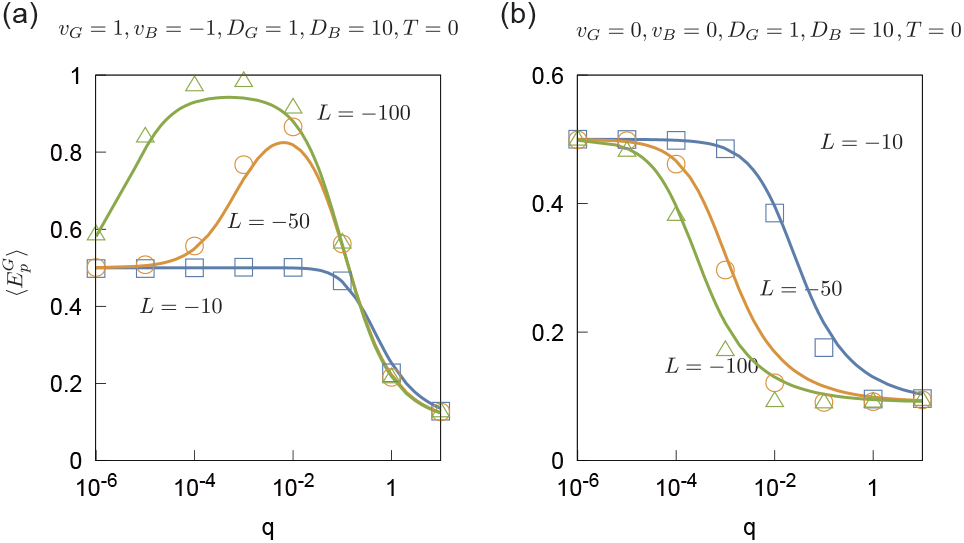
Theoretically-predicted and numerically-evaluated decision-bias induced by the random decision switching. **(a)** For the emergence of globally-persistent decision-bias on GD, |*H*− *L* |≫10 and moderate *q* are necessary. For *q >* 0.1, the bias toward BD emerges. The lines are theoretically obtained and the points are obtained by the numerical simulation. **(b)** For *v*_*G*_ = *v*_*B*_ = 0, only the decision-bias toward BD, i.e., an option having the larger deviation, exists. Theoretical predictions were conducted using Mathematica with 1500 significant digits. The discrepancy between the theoretical predictions and the numerical simulations may be due to the fact that the theoretical equation includes the term *e*^*−H*+*L*^), which yields an extremely small value with *H* − *L >* 50. Each point was computed by running 2 × 10^5^ numerical simulations.

Although Eq. 21 and 23 are theoretically exact, due to the numerical instability of higher-order modes, it can be numerically-evaluated only with the moderate parameter region such as |*H* –*L*| *<* 10, and *q* ≪1. The numerical instability is pronounced when the initial probability distribution involves discontinuity as Dirac’s delta function or the uniform distributions, because representing the discontinuous initial distributions requires many higher-order modes. Thus, we need to consider an alternative approach to estimate the total decision bias. In particular, we consider the total probability of escaping as GD, 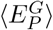, which is described as

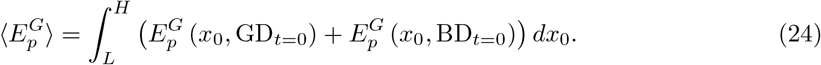

Hereafter, we describe 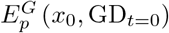 as 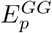and 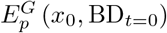 as 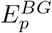for simplicity. 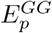 and 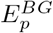 are computed by solving KBE 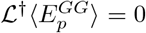 and 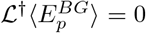. Here, L† is exactly the same with Eq. 17, but the boundary condition is different: 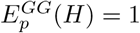 and 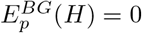 (i.e., a particle started with GD/BD and *x*_0_ = *H* is immediately absorbed as GD/BD), 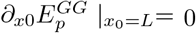 and 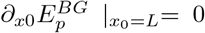. Fig. 5-(d) shows that globally-persistent decision-bias emerges for | *H* − *L* |≫ 10. This implies that separating *P*_*G*_ and *P*_*B*_ requires enough space depending on *v*_*G*_, *v*_*B*_, *D*_*G*_, *D*_*B*_, and *q*. Understanding the minimum size of the space will be the future direction.

For such globally assymetric regime, *q <* 0.1 leads to the decision-bias to GD and *q >* 0.1 leads the decision-bias to BD. In *q* ≫ 0, the diffusion force is dominant because the drifting forces cancel each other out as *v*_*G*_ + *v*_*B*_ ∼0. Therefore, in this regime, the state of the thermal bath with the larger noise has a higher chance of reaching the absorbing boundary, and the decision is biased towards BD. Furthermore, it is noteworthy that the bias toward BD is noise-induced metastable state.

## 6 Discussion

### 6.1 Multi-layered decision biasing and flipping

In the general framework of decision-making, the outcome of a decision is subjectively evaluated by a utility function [33], and the subjective value score *U*_*n*_ influences the subsequent decision-making. In contrast, our embodied decision-making system assumes that the subjective value space, *U*, is mapped onto the IBA parameter space, *X*, using an operator 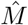as 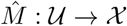. Here, *X* has different functional phases, Sw and Loc, which generate information asymmetry, and this asymmetry induces decision biases in a self-organizing manner.

The decision-biasing and -flipping mechanisms are multilayered. The Sw region induces a first-passage decision bias, while the Loc region rectifies this bias. By changing the sign of 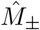, the overall decision bias is flipped (we use the notation 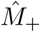 when *U*_*n*_ *>* 0 is mapped to *x >* 0, and 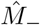 when *U*_*n*_ *>* 0 is mapped to *x <* 0). In addition, the random rate *q* can be controlled by adjusting the rhythmic activity and decision sampling timing. Changing the random rate *q* can flip the first-passage bias. Here, we outline this mechanism (see also Fig. 7).

**Figure 7:**
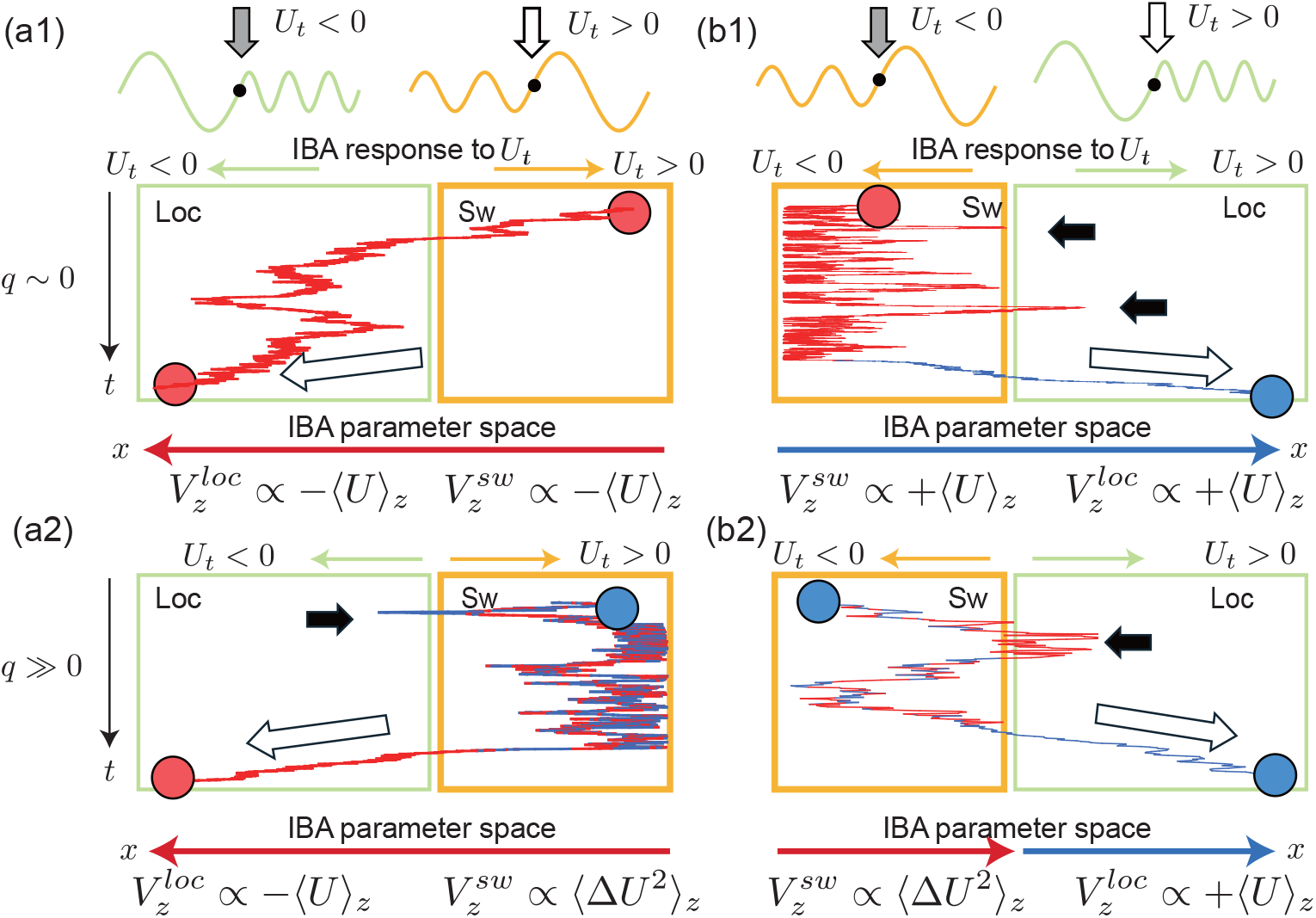
Understanding the multilayered decision bias created by rhythmic coupling of an internal organ and the brain. The IBA in response to the subjectie value representation *U*_*t*_ provides an additional degree of freedom to construct a body-guided value space. Whether the decision outcome *U*_*t*_ *>* 0 contributes to the decision locking is determined by how IBA is updated in response to *U*_*t*_. On the left panels **(a1, a2)**, *U*_*t*_ *>* 0 is mapped to IBA responses toward the Sw region. Therefore, only decision options with an expected value ⟨*U* ⟩ *<* 0 are stably locked (options with ⟨*U* ⟩ *>* 0 are rejected in the Loc region, as shown by the bold arrow). Similarly, on the right panels **(b1, b2)**, *U*_*t*_ *>* 0 is mapped to IBA responses toward Loc regions. Therefore, only decision options having ⟨*U* ⟩ *>* 0 are stably locked. For *q* ∼ 0, the value function with respect to decision option *z, V*_*z*_, is the same in the Loc and Sw regions. However, for *q* ≫ 0, the value functions in the Loc and Sw regions (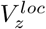 and 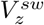, respectively) differ. In the Sw regions with *q* ≫ 0, the drifting terms of options are cancels out each other, and only diffusion terms are important. Therefore, 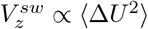 . If an option which reached the Loc region does not have the drifting term matching the value function *V* ^*loc*^, such option is rejected in the Loc region, as shown by the bold arrows. Red and blue arrows denote the biases toward BD and GD, respectively.

**Figure 8:**
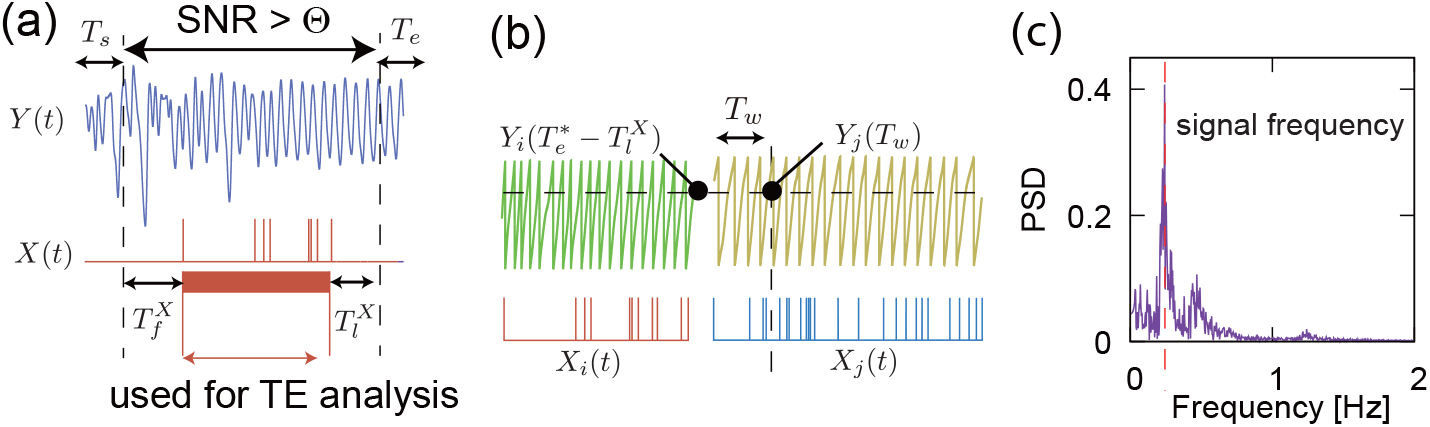
Preparing data for TE and MCMC analyses. **(a)** The cropping procedure for the pair of source process *Y* (*t*) and target process *X*(*t*). We have searched for the time window [*T*_*s*_, *T*_*e*_] that satisfies SNR *>* Θ or otherwise maximizes the SNR of *Y* (*t*). The pair (*X*(*t*), *Y* (*t*)) is clipped by the time window 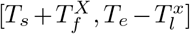, where 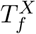 and 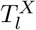 are the time to the first and last event spikes, respectively. **(B)** (*X*_*i*_(*t*), *Y*_*i*_(*t*)) and (*X*_*j*_(*t*), *Y*_*j*_(*t*)) of the *i*th and *j*th participant are connected at the time *t* = *T*_*w*_ which gives *d* 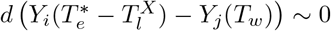. **(C)** A brief description of how to determine the signal frequency of *Y* (*t*) by using the FFT; we assumed the signal frequency *ω*_*s*_ has the maximum power spectrum density (PSD) *P* (*ω*_*s*_).

#### Loc region as a rectifier

If the decision outcome *U*_*t*_ *>* 0 is mapped by 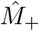 so that the IBA state moves away from the Sw region (i.e., toward *x >* 0), then, the Loc region stabilizes a decision option *z* having⟨ *U* ⟩_*z*_ *>* 0. With this mapping function 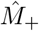, an option *z* having ⟨*U* ⟩_*z*_ *<* 0 are rejected as shown by the black arrows in the Fig. 7-(b, c, d). On the other hand, if the decision outcome *U*_*t*_ *<* 0 is mapped by 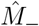 so that the IBA state moves away from the Sw region, a decision option *z* having ⟨*U* ⟩_*z*_ *<* 0, is stabilized in the Loc region. Thus, the Loc region acts as a rectifier, allowing only options with the “correct” expected value to persist where the correct direction is determined by the mapping function 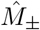.

#### First-passage bias in the Sw region

Independently to the Loc region, the Sw region alone can form first-passage decision biases. With *q* ∼ 0, the first-passage decision bias is formed based on the expected value structure of options ⟨*U*⟩ _*z*_, similarly to the Loc region. Those options which move *x* toward *x* = *L* are trapped within the Sw region, and do not reach the Loc region. When *q* ≫ 0, the decision output often changes a nd the drifting force of x asymptotically approaches the sum of the expected option values, i.e., Σ_*z*_⟨*U* ⟩_*z*_. (In most setups in Fig. 3 –5, *v*_*G*_ + *v*_*B*_ = 0, so drifting terms cancels each other out perfectly). In this case, the effective driving force of *x* is provided by the diffusive terms of the option values ∝ Σ _*z*_ ⟨Δ*U* ^2^⟩ _*z*_, and decision output is biased toward options with larger variance (In our experimental setupt, BD). However, if the options selelcted by the first-passage bias does not have the “correct” expected value ⟨*U* ⟩_*z*_, such options are rejected by the Loc region. Consequently, only those options with a value of ⟨*U*⟩ _*z*_ that matches the direction of the rectifier in the Loc region can persist.

#### Body-guided decision-making overrides the subjective value representation

In the framework of inverse reinforcement learning [64], the probability of taking a certain behavior *z, P*_*z*_ and the value function *V*_*z*_ are connected by the Boltzman-like distribution as *P*_*z*_ = exp (*V*_*z*_) */Σ* _*z*_ exp (*V*_*z*_), where the value function is estimated as *V*_*z*_ ∝ log *P*_*z*_. Therefore, the overall process of body-guided decision-making is a process which overrides *U*_*n*_, the representation of subjective value of each decision outcome, to the exteranlly observable value function *V*_*z*_ of the decision option *z*, via the decision bias *P*_*z*_. In addition, as shown in Fig. 7, for *q* ∼ 0, the value function of the option *z* in the Sw region is 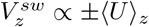 (an option with the correct expected value is valuable), and for 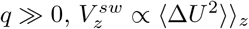 (an option with large score deviation is valuable).

### 6.2 Emergent arbitrariness and agent’s free discretion

To the best of our knowledge and consideration, there are no biological or physical laws that determine whether the mapping function should be 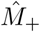 or 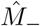. Regarding emotions in animals and humans, it is suggested that a “concept of a particular emotion” (e.g., fear is associated with a fast heartbeat) plays a crucial role in triggering physiological reactions based on the subjective evaluation of a situation. This concept is either genetically acquired or culturally constructed [8, 4]. However, the ability to shape emotional responses through “culturally constructed emotion concepts” [4] suggests that emotions imply the presence of “meta-level decision-making” or “free discretion” and “free will”. Within the original SMH framework, the ventromedial prefrontal cortex (vmPFC) [14, 15] must be responsible for coordinating physiological changes based on subjective evaluations onto internal organs based on allostatic functions [52].

In fact, we had expected that human rationality could be fully explained deterministically and causally through the rhythmic coupling of the internal organs and the brain. We had expected that decision-making bias could be uniquely determined by the presence of rhythmic IBA. However, we found that there remains an element of arbitrariness and agent’s discretion. The most important factor determining the bias was the sign of the mapping function 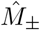. We hope that this remaining or emergent arbitrariness in the body-guided decision-making is essential to building and realizing intersubjectivity, as discussed in the following section.

### 6.3 Intersubjectivity beyond subjectivity

Fig. 4-(a) show the surprising results that reward-modulated Hebbian learning, which we believe is a simple, neural-level implementation of the value-based reasoning process, corresponding to Bayesian computation within a brain [39, 44, 36], cannot produce rational decision-making when the internal evaluative utility function is even slightly biased toward underestimating risk. This result highlights a serious issue: while a learning process based on an subjective evaluation function is reasonable from the system’s viewpoint, it is considered irrational by an outside observer. Therefore, rationality cannot be achieved by the local learning rule in the brain alone.

To correct distorted subjective evaluations and establish a “rational” decision-making process aligned with external consensus, one needs another mechanism which can share information with an “external” existence. (This external existence could be another person or society, for example). Our results suggest that the mapping subjective evaluations onto internal organs by the function 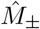 can sufficiently correct these distorted evaluations, yielding self-organization of rationality that aligns with external consensus.

The subjective representation of the decision outcome, *U*_*n*_, is essentially neither observable nor shareable with others. However, the behavioral bias *P*_*z*_ is observable, so the value function of the option *z* can easily be accessed as *V*_*z*_ ∝ *logP*_*z*_. Furthermore, behavioral bias can be controlled by changing how bodily activity is updated. This mechanism provides an opportunity to share and coordinate the value function *V*_*z*_ with others to create a coordinated intersubjective world.

### 6.4 Impact of rhythmic IBA

Fig. 3-(d) demonstrates that the decision bias can be formed only by updating the average level of IBA, *g*_*b*_, without changing the amplitude *g*_*g*_, Δ*g*_*g*_ = 0. Furthermore, large decision bias can be formed even when there is no periodic activity, i.e., *g*_*g*_ = 0. This occurs because the decision outcomes inovolve stochastic variables,therby inducing spontaneous decision switching in the Sw at very small switching rate *q*∼ 0. Does this mean that rhythmic IBAs are not necessary for creating decision-making biases? This issue should be considered from the following two perspectives.

#### The rhythmic IBA explains human decision-making strategy

The rhythmic IBA induces spontaneous decision switching independently the randomness induced by the decision outcome. This enables controlling *q*, the random decision switching rate. For example, if IBA is in *f* [Hz] and the decision-making is executed after *T* [s] such that *T* is sufficiently larger than 1*/f* [s], then, the next decision output will be different from the current one. In contrast, if *T* is less than 1*/f* [s], the decision output will be the same as the current one. Thus, *q* is controlled via the timing of decision execution, *T* . The controllability of *q* is one of the distinct advantages of rhythmic IBA.

Furthermore, if there is no rhythmic IBA, meaning that *q* is not controllable and only *q*∼ 0 is available, the overarching decision strategy of initially prioritizing score variance (BD bias) and switching to options with a higher expected value (GD bias) when encountering a large negative output disappears. This overarching decision behavior, also shown in Fig. 7-(c, d), is often framed as reversal learning and is often seen in human IGT experiments.

#### Biological indivisibility of the rhythmic amplitude and average activity

Recent studies report that LC activity is strongly enhanced by the reward prediction error which may be primarily computed by cortical neurons [30, 31]. However, many peripheral, rythmic activities such as cardiovascular activity, pulmonary activity, partial pressure of carbon dioxide (pCO_2_), licking (the movement of a tongue), influence LC activity [54]. Furthermore, strong burst firing of LC neurons inevitably leads to the strong autoinhibition mediated by *α*_2_ receptors [3], leading the emergence of “one cycle” of rhythmic activity of LC bursting neurons. Therefore, even when discussing the LC average activity, it seems impossible to separate biological rhythmic activity from average activity. We hope the results shown in Fig. 3-(d) are considered as being biologically plausible. In humans, a decrease in *g*_*g*_ and an increase in *g*_*b*_ correspond to shallow breathing, which leads to high pCO_2_, inducing high LC activity (and hence high neuronal nonlinearity *g*). Conversely, an increase in *g*_*g*_ and a decrease in *g*_*b*_ correspond to deep breathing, which leads to low pCO_2_, inducing low LC activity (and hence low neuronal nonlinearity *g*).

#### Strong consistency with biological data

It should be noted that our results are strongly consistent with recent biological evidence indicating that silencing the locus coeruleus results in changes in decision-making, and when rewards are obtained unpredictably based on a particular decision, activation of the locus coeruleus reinforces that decision [53]. In addition, we point out that our system can reproduce the win-stay/lose-shift strategy, which better explains human behavior in IGT than reinforcement learning [60]. In summary, these results and evidence suggest that human decision-making is supported by IBA.

### 6.5 Homeostasis, rhythmic activity, and decision-making

Although our framework does not explicitly incorporate homeostasis, we assume that periodic IBA captures its essence. Many human internal organs exhibit periodic dynamics. For example, gastrointestinal organs operate at frequencies of around 0.01–0.1 Hz, respiratory activity occurs at frequencies of around 0.1–0.5 Hz, and cardiac activity occurs at frequencies of around 1–3 Hz [21]. These periodicities arise from the need to balance nutrient storage with behavioral freedom. In other words, animals store nutrients so that they can move freely in their environment. Otherwise, they would always need to be in contact with a source of nutrients, which would restrict their behaviour. Thus, considering the rhythmic IBA involves the issue of homeostasis. A weakness of our current framework is that we did not consider the boundary of the allowable range of homeostasis; we assumed that the IBA itself was not limited to a certain range, although we did assume that the neuromodulator intensity *g*(*t*) *>* 0. Investigating how the behavior of the system changes when there are upper or lower limits on the IBA is a futur work.

Recently, the concept of reward in animals has been updated; it is shown that the primary reward capable of directly stimulating the brain’s reward system is the digested and absorbed nutrient itself, meaning that taste and smell do not qualify as primary rewards. Instead, taste and smell are considered as proxy rewards that serve to sustain behavior until nutrients are digested and absorbed [58]. Our results have elucidated how neural decision-making output is switched or locked by the activity of internal organs. Therefore, the Sw-Loc mechanism may also be involved in the proxy reward mechanism that helps sustain current behavior.

### 6.6 Body-guided decision-making as embodied Bayesian-like update

Our results reveals additional perspective of decision-making. Since the Langevin and Fokker-Planck equations are mathematically equivalent, sequentially updating IBA’s parameters *x* based on stochastic decision outcomes is equivalent to updating the probability *P*_*z*_(*x, t*) which represents the belief that the probability that an option *z* (*z* is GD or BD) is selected at the current state *x* and the time *t*. Consequently, body-guided decision-making tasks can be viewed as performing Bayesian-like updating of belief *P*_*z*_(*x*), based on the observed data. In the latter Bayesian framework, the current state *x* and the option *z* at the time *t* are considered to be sampled from *P*_*G,B*_(*x, t*), and the magnitude of the belief *P*_*z*_ at *x* = *H* − *ϵ* (*ϵ* → 0) determines which option *z* is likely to be biased. Importantly, the internal bodily parameter *x* is merely mediates the belief *P*_*z*_; it does not directly represent the value *V*_*z*_ of the option *z*. Rather, *V*_*z*_ emerges indirectly through the formation of a body-guided behavioral bias, as illustrated in Fig. 7. Our results fully account for the recent finding that the message conveyed by a physiological state shifts depending on the situation [20]. Thus, our results offer a thorough explanation of the somatic marker hypothesis, suggesting that decision-making occurs at the intersection of internal and external information.

## 7 Conclusion

The present study showed that perceptual decisions in human binocular rivalry are influenced by the activity of internal organs. Through computational and mathematical analysis, we showed that internal organs can bifurcate the decision-making output in a brain to lock and switch modes. Through computational simulations of the IGT, we demonstrated that decision-making agents who receive rhythmic information from internal organs can make rational decisions without knowing or estimating the value of decision outcomes. We demonstrated that reward-based Hebbian learning can be easily corrupted by biased subjective reasoning and evaluation systems that underestimate risk. However, coupling the brain with the internal organs can enable rational decision-making. We revealed a core mechanism of SMH: the rhythmic activity of the internal organs bridges subjective reasoning and intersubjective rationality.

## Acknowledgement

This study was partially funded by Toyota Central R&D Labs., Inc., Chair for Frontier AI Education of the Next Generation AI Research Center, The University of Tokyo, and JSPS KAKENHI Grant Number JP 25K24741. Our theoretical treatment described in Sec. 5 were conducted with the help of ChatGPT (GPT-5.2 later) and Gemini 3.5 Thinking mode and verified by authors using Mathematica.

## A Experimental setup of binocular rivalry

The binocular rivalry experiment described in this paper was approved by the ethics committees of our institute. All participant for the binocular rivalry experiments signed an informed consent statement. Visual stimuli with different stripe angles are presented to the left and right eyes of a participant, and a participant is asked to press either the left or right button, or both buttons, when he/she perceives a visual image presented to the left or right eye, or a mixed image. The time of the perceptual alternation *e*_*p*_(*t*) was recorded.

The experiment started 3 minutes after the first training session. In the first session (6 minutes), a participant is asked to continue breathing at normal speed. After the 3 minutes of rest, a participant is asked to continue breathing at a fast rate. In addition, after the 3 minutes of rest, a participant is asked to breathe at a slow rate. The actual breathing rate is at the discretion of the participant, as we did not use a metronome to dictate the breathing rate. This is because the natural breathing rate must vary from person to person, and breathing at an unnatural rate may have unpredictable effects on the mental state or perceptual decision-making process.

The change in a participant’s chest volume *v*_0_(*t*) was measured at 500 Hz using a belt sensor; DC respiratory sensor, manufactured by Miyuki Giken, Tokyo, and cardiac activity was measured by an electrocardiogram manufactured by METS, Inc. at 500 Hz. Pupil diameter was measured with an HTC VIVE Pro Eye eye tracking device at 100 Hz.

### Preprocessing data for TE and MCMC analyses

In TE analysis, we need paired time series data of the source process *Y* (*t*) and the target process *X*(*t*). In the following, we describe the preprocessing procedure for *X*(*t*) and *Y* (*t*). For the MCMC analysis, we used the preprocessed data of the source *Y* (*t*) = *θ*(*t*) and *X*(*t*) = *e*_*p*_(*t*) described below.

#### Bandpass filters

We applied the bandpass filter of [0.03, 1.0] Hz to the value of a belt sensor *v*_0_(*t*) to obtain *v*(*t*). The phase *θ*(*t*) of the chest motion is calculated from the Hilbert transform of the bandpassed signal *v*(*t*) as *θ*(*t*) = *H*(*v*) + 1*/*2*π*. Note that we add a phase shift of 1*/*2*π* so that *θ*(*t*) = *π/*2 at the peak of *v*(*t*). The cardiac R-wave *R*(*t*) was computed using the wavelet transform [26]. The bandpass filter of [0.1, 3] Hz is applied to the pupil diameter and we obtain *d*_*p*_(*t*) : [0, 1] by normalizing the bandpass signal. Blink times *B*_*l*_(*t*) were estimated by detecting the sudden and large fluctuation of the pupil diameter.

#### Cropping of paired processes of a source Y(t) and a target X(t) for artifact and transient removal

The TE measure assumes that the source process *Y* (*t*) and the target process *X*(*t*) are stationary [50]. However, human physiological data, especially respiration data *v*(*t*), contain many artifacts, such as voluntary changes in respiration rate, belt sensor position shifts, and these artifacts create transient and nonstationary respiration data. To crop the stationary region of the paired data of (*X*(*t*), *Y* (*t*)), we used the time window [*T*_*s*_, *T*_*e*_] and calculated the signal-to-noise ratio (SNR) of *Y* (*t*) within the time [*T*_*s*_, *T*_*e*_]. Note that the SNR will be higher if *Y* (*t*) is stationary and lower if *Y* (*t*) has non-stationary characteristics, such as changes in respiratory rhythm or other artifacts. Furthermore, we assumed that if *Y* (*t*) is stationary, *X*(*t*) is also stationary. (To obtain enough data-length, we assumed *T*_*s*_ and *T*_*e*_ is at most 100 [s]).

The SNR is calculated as follows: We computed the power spectrum 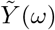 of *Y* (*t*) using the Fast Fourier Transform (FFT), where *ω* is the frequency. We assumed that *ω*_*s*_, which gives the maximum 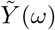, is the signal frequency of *Y* (*t*). The power spectrum density *P* at the discrete frequency step *k* used in the FFT is calculated as 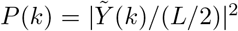 where *L* is the length of the data used in the FFT. Then the SNR is 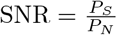, where 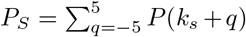 is the power spectrum density of the signal frequency, and *P*_*N*_ is the sum of the power spectrum densities other than the signal frequency, i.e., *P*_*N*_ = *P* (*k*) −*P*_*S*_.

We searched for the time window 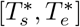 that satisfies SNR *>* Θ or otherwise gives the maximum SNR, and the paired data of the source *Y* (*t*) and the target *X*(*t*) are cropped by the time window 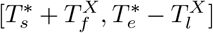, where 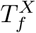and 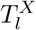 indicate the time to the first and last event spike of *X*(*t*) after 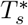and before 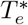, respectively, as shown in Fig. 9. Note that 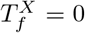 and 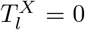 if *X*(*t*) is a nonpoint process.

**Figure 9:**
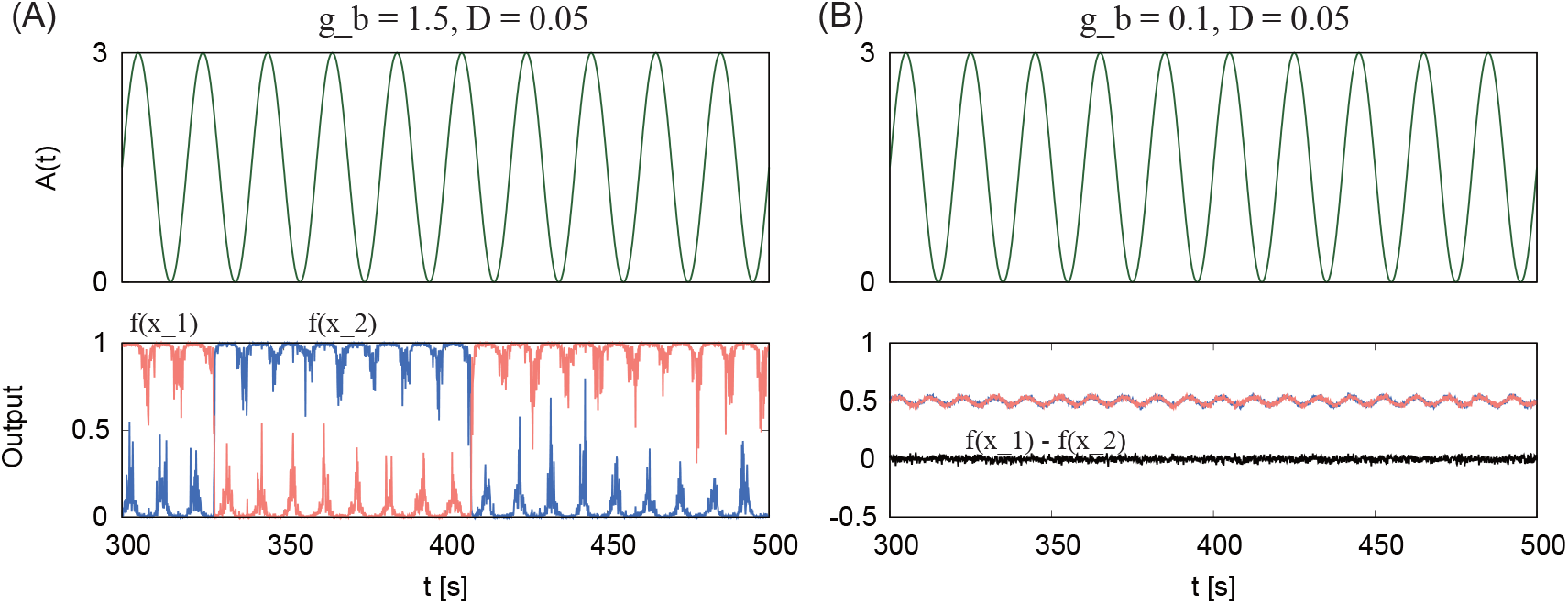
WTA behavior in response to the additive input signal *A*(*t*) with moderate activation gain *g*_*b*_ = 1.5. (**A**), and small activation gain *g*_*b*_ = 0.1 (B).It is shown that with moderate *g*_*b*_, WTA exhibits bistable behavior and stochastic switching behavior based on the input *A*(*t*). The bistability is lost for small *g*_*b*_, and in this parameter regime, *f* (*x*_1_) − *f* (*x*_2_) ∼0. We used *D* = 0.05.

**Figure 10:**
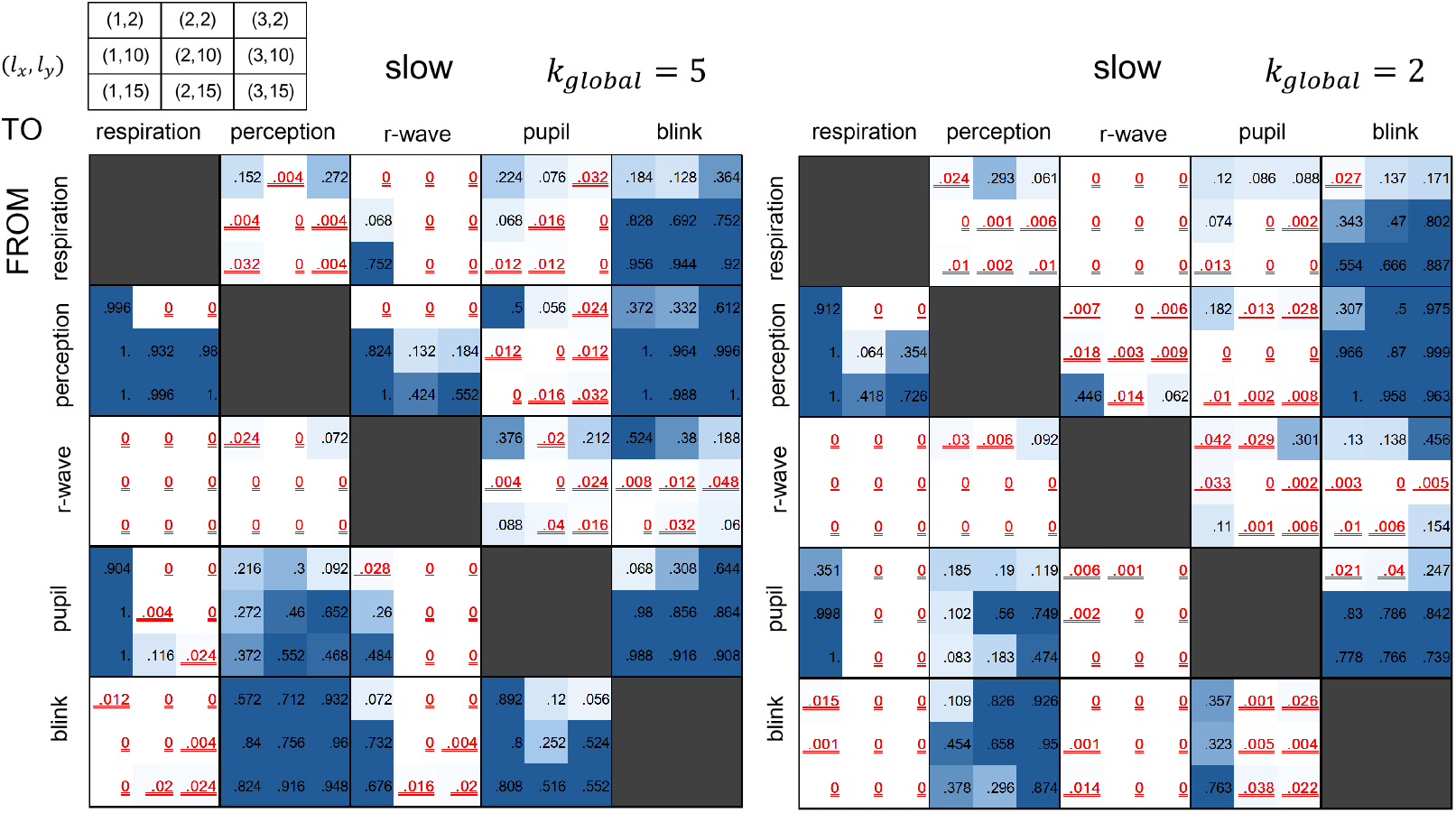
The heatmap shows the p-values obtained by means of the permutation test using transfer entropy (TE) between the respiration phase *θ*, the timing of the perception alternation *e*_*p*_, the R-wave, *R*, the normalized pupil diameter *d*_*p*_, and the blink timing *B*_*l*_ in the slow respiration rate. We used *k*_*global*_ = 5 and *k*_*global*_ = 2 to calculate TE. Furthermore, we used *k*_*perm*_ = 100 for *k*_*global*_ = 2. The 3 ×3 submatrix denotes the embedding dimension *l*_*x*_, *l*_*y*_ used to compute TE from *y* → *x* (*y* and *x* are the source and target processes, respectively). The p-values which satisfy the condition *<* 0.05 are indicated by blue fonts with a double underline.

When the respiration phase *θ* is a source *Y* (*t*), we calculated the SNR using the respiration data *v*(*t*), because the phase *θ* loses the information of the signal amplitude (and therefore the SNR using *θ*(*t*) has less sensitivity to detect the transient process). Furthermore, when *Y* (*t*) = *θ*(*t*) and the maximum SNR is less than 0.3, we did not use the whole *v*(*t*) because it means that *v*(*t*) is not stationary. Similarly, we did not use some data with a significantly low number of 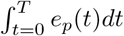 (where *T* = 6 min is the session duration), e.g, only one perceptual alternation during a 6-min session, because it seems that the recording of *e*_*p*_(*t*) failed; we did not use the data *e*_*p*_ if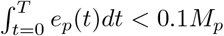, where *M*_p_ is the average number of perceptual alternations of all participants. Consequently, we collected the pair data of (*v*(*t*), *e*_*p*_(*t*)) from 18 participants for normal breathing rate, 11 participants for fast breathing, and 17 participants for slow breathing out of 30 participants (30 male, mean age is 25.5 with standard deviation 2.8).

**Figure 11:**
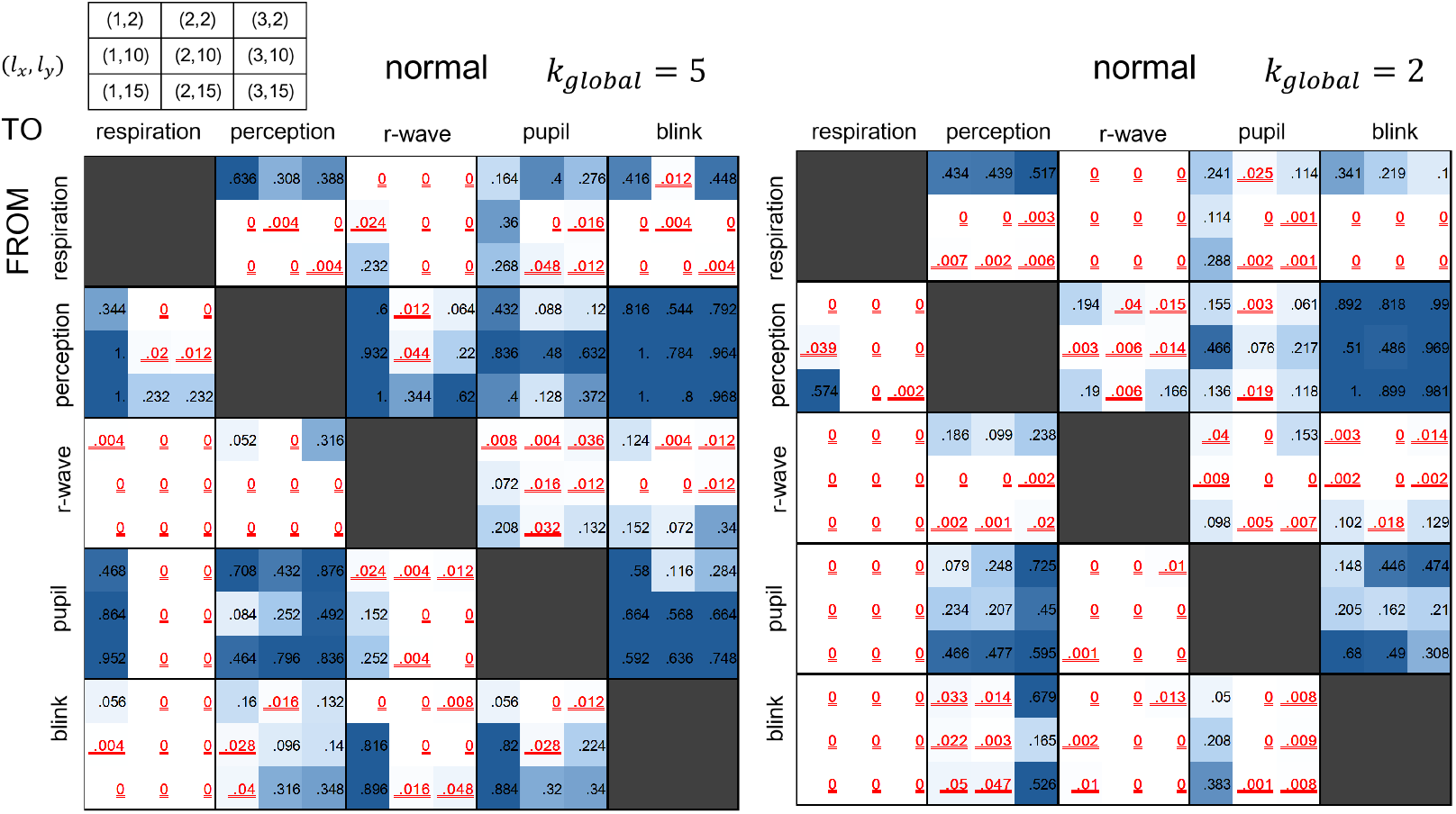
The heatmap shows the p-values obtained by means of the permutation test using transfer entropy (TE) between the respiration phase *θ*, the timing of the perception alternation *e*_*p*_, the R-wave, *R*, the normalized pupil diameter *d*_*p*_, and the blink timing *B*_*l*_ in the normal respiration rate. We used *k*_*global*_ = 5 and *k*_*global*_ = 2 to calculate TE. Furthermore, we used *k*_*perm*_ = 100 for *k*_*global*_ = 2. The 3 × 3 submatrix denotes the embedding dimension *l*_*x*_, *l*_*y*_ used to compute TE from *y*→ *x* (*y* and *x* are the source and target processes, respectively). The p-values which satisfy the condition *<* 0.05 are indicated by blue fonts with a double underline.

**Figure 12:**
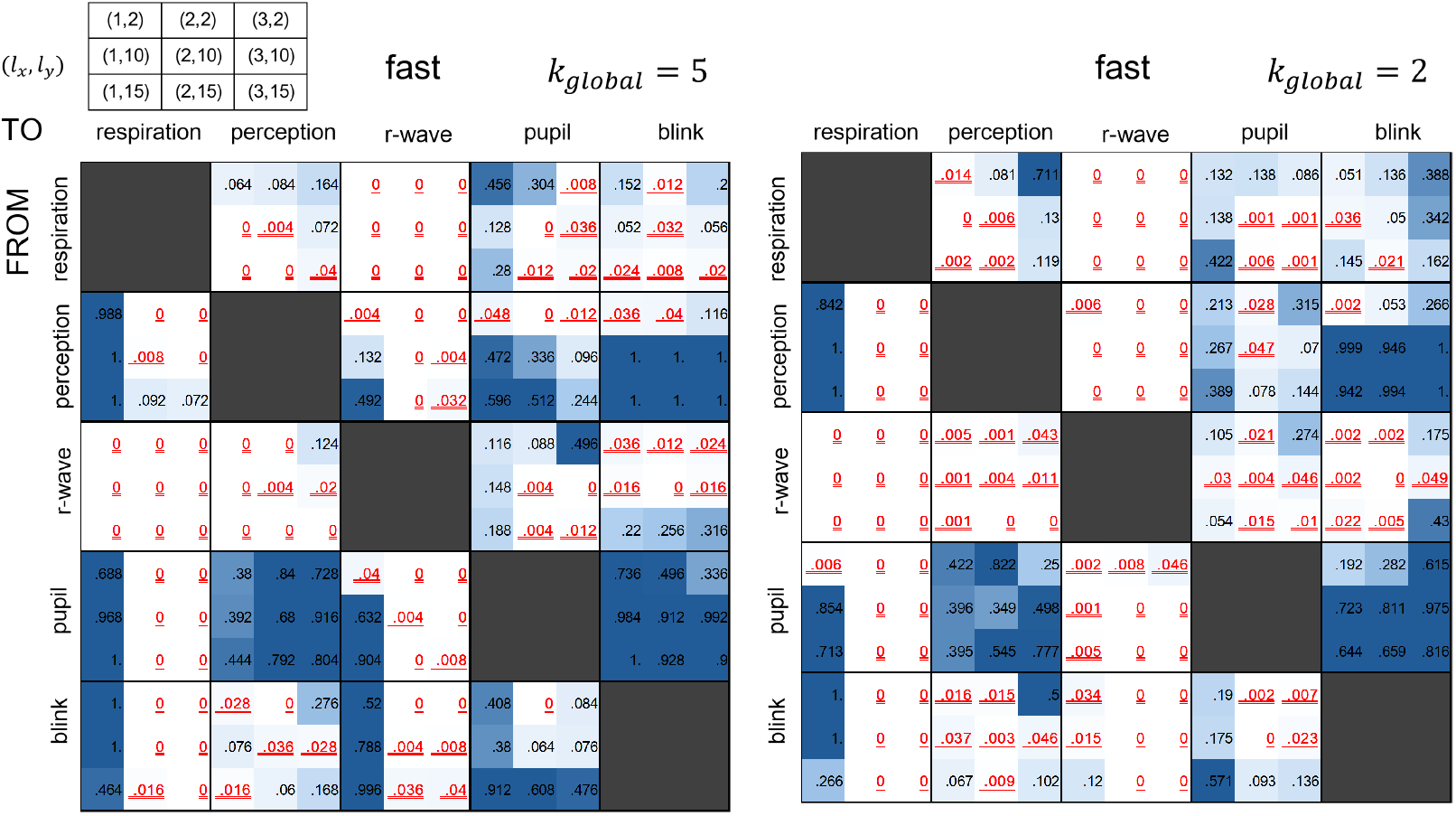
The heatmap shows the p-values obtained by means of the permutation test using transfer entropy (TE) between the respiration phase *θ*, the timing of the perception alternation *e*_*p*_, the R-wave, *R*, the normalized pupil diameter *d*_*p*_, and the blink timing *B*_*l*_ in the fast respiration rate. We used *k*_*global*_ = 5 and *k*_*global*_ = 2 to calculate TE. Furthermore, we used *k*_*perm*_ = 100 for *k*_*global*_ = 2. The 3 ×3 submatrix denotes the embedding dimension *l*_*x*_, *l*_*y*_ used to compute TE from *y* →*x* (*y* and *x* are the source and target processes, respectively). The p-values which satisfy the condition *<* 0.05 are indicated by blue fonts with a double underline.

#### Merging the cropped data from multiple participants

The cropped stationary data from different participants are merged into a time series data. To merge *Z*_*i*_ = (*X*_*i*_, *Y*_*i*_) and *Z*_*j*_ = (*X*_*j*_, *Y*_*j*_), we searched for *T*_*w*_ that satisfies 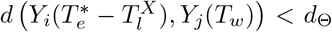, where 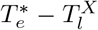 is the last time stamp of (*X*_*i*_, *Y*_*i*_), *d*(*x, y*) is a Euclidean distance, and *d*_Θ_ = 0.05. It should be noted that this merging strategy does not break the structure of the flow of information from *Y* to *X*.

If both the source and target processes are nonpoint processes, we simply computed the Euclidean distance *d*(*Z*_*i*_(*T*_*e*_), *Z*_*j*_(*T*_*w*_)) and searched for the minimum *T*_*w*_ that gave *d <* 0.05. If both the source and destination processes are point processes, we simply merged *Z*_*i*_ and *Z*_*j*_ at *T*_*w*_ = 0.

### Converting a nonpoint process to a point process

We computed the TE between respiration phase *θ*(*t*), cardiac R-wave *R*(*t*), pupil diameter *d*_*p*_(*t*) and blinking timing *B*_*l*_(*t*) and perceptual alternation event *e*_*p*_(*t*) by using the computational method provided by Ref. [50]. This method assumes that both source and target signals, *Y* and *X* are point-process and have the continuous time representation; 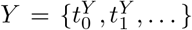, and 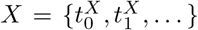, where 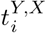 is the timestamp of the event occurrence.

It is important to note that *R*(*t*), *B*_*l*_(*t*) and *e*_*p*_(*t*) are point processes and *θ*(*t*) and *d*_*p*_(*t*) are continuous processes. Therefore, it is necessary to convert *θ*(*t*) and *d*_*p*_(*t*) to point processes. To convert the *q*(*t*) of a continuous process to a point process time series *Q*^*p*^(*t*), the variable space is divided into 10 subspaces [Ω_0_, Ω_1_, … Ω_9_]. If the continuous signal *q*(*t*) moves from the current subspace to another subspace at time *t*^*q*^, the time stamp *t*^*q*^ is appended to the time series *Q*^*P*^ (*t*). This results in the following representation: 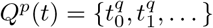.

Note that the TE value depends on the seed for the random number generator. Therefore, we calculated the TE value using 10 different random seeds. In addition, we generated 250 surrogate data for each seed. We calculated *p* value using 5 original TE values and 1250 surrogate TE values; *p* = *N*_*s*_*/N*_*total*_, where *N*_*s*_ is the number of surrogate data that have a higher TE value than the original data, and *N*_*total*_ = 1250.

#### The parameters used to calculate TE

The parameters used to calculate TE are tabulated below. For detailed explanation of the parameters, see Refs. [50]. It should be noted that the respiratory phase *θ* has a significant information flow with *l*_*y*_ ≥ 10 with *x* = 2, 3. This implies that one cycle of breathing has a significant contribution to two and three perceptual switching events ^1^.

**Table 1:** The parameters used to calculate TE. For Fig. 1 we used *k*_*global*_ = 5 and *k*_*perm*_ = 50. The results of the permutation test using TE with *k*_*global*_ = 2, *k*_*perm*_ = 100, and *I*_*x*_ = 1 are shown in the supplementary Fig. 10–12.

| Parameter | Description | Value |
| --- | --- | --- |
| $I_x$ | Number of interspike intervals in target history embeddings | 1, 2, 3 |
| $I_y$ | Number of interspike intervals in source history embeddings | 2, 10 |
| $k_{global}$ | Initial number of nearest neighbors to determine the radius of the ε-ball | 2, 5 |
| $N_X$ | Number of target events in the data from a participant | varied |
| $N_U/N_X$ | Ratio of random samples used to estimate the unconditional probability density of the target process | 10, 1 |
| $N_{surrogates}$ | Number of surrogates data | 250 |
| $k_{perm}$ | number of nearest neighbours used in the local permutation tests | 100, 50 |
| $D_k$ | Small noise intensity added to construct the embedding vector $(x, y)$ | $10^{-8}$ |

#### The parameter sensitivity of TE

The results of the permutation test using TE between point processes [50] have a weak sensitivity to the parameter especially *k*_*global*_ which is the parameter to determine the radius of the *ϵ*-ball; to explain it in a very simple way, the TE algorithm counts the number of points inside the *ϵ*-ball. Note that this algorithm assumes that the data points are uniformly distributed within the *ϵ*-ball. Therefore using too large value of *k*_*global*_ will result in incorrect TE estimation if the data length is not long enough.

The supplementary Fig. 10–12 shows the differences of the p-values computed with *k*_*global*_ = 2 and 5. It is shown that *k*_*global*_ = 2 marks the information flows as significant in more combinations of the embedding dimensions (*I*_*x*_, *I*_*y*_) than *k*_*global*_ = 5. Ref. [50] argues that both *k*_*global*_ = 2 and 5 have sufficient accuracy to detect and reject the significant and insignificant information flows if the data length is long enough. The difference in *p* values for different (*I*_*x*_, *I*_*y*_) induced by *k*_*global*_ in our analysis of the human data may reflect the data length limitation. Note that the different *p* values induced by *k*_*global*_ do not affect our main results that there are significant mutual information flows between perceptual alternation and interoception.

## B The simplest neural network model for perceptual and behavioral decision-making

The computational model for perceptual and behavioral decision-making shown in Fig. 2 is described as

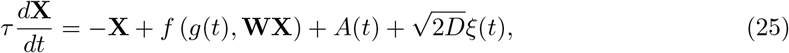

where 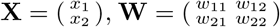, and 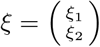, and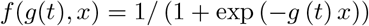. Note that *g*(*t*) is the activation function of a neuron, reflecting the effect of the neuromodulator (described in further detail below). Additionally, the condition *g*(*t*) *>* 0 must be satisfied for the neurons to retain bistability and winner-take-all (WTA) capability. *x*_*i*_ (where the indices *i* and *j* are 1 or 2) denotes the membrane state of *i*th neuron, *f* (*x*) denotes the sigmoidal activation function of a neuron which roughly corresponds to the firing rate of a neuron, *w*_*ii*_ and *w*_*ij*_ represents the weight of self-excitatory and mutual-inhibitory synaptic connections, respectively. 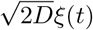 denotes the Gaussian noise with the standard deviation of *D*. In Eq. 2 in the main text, we use the notations *w*_*exc*_ and *w*_*ii*_ to express the excitatory self coupling *w*_11_ or *w*_22_. Similarly, we use *w*_*inh*_ and *w*_*ij*_ to express the mutual inhibitory coupling *w*_12_ or *w*_21_. Furthermore, we use the parameters *w*_*exc*_ = *w*_*ii*_ = 3*/*2 and *w*_*inh*_ = *w*_*ij*_ = −9*/*2.

### Numerical simulations of WTA

The data shown in Fig. 3 were obtained by using the results of 50 trials of a 10^5^ s numerical simulation. The stepsize of a numerical simulation was 1 [ms], and a different random seed was used in each simulation trial. The human perceptual state has at least three states: left-image dominant (L), right-image dominant (R), or mixed (LR). The transition between these three states in numerical simulations using a WTA circuit can be represented by using the variable of *F* (*x*_1_, *x*_2_) = *f* (*x*_1_) − *f* (*x*_2_). However, we have defined only two states L: *F* (*x*_1_, *x*_2_) *>* 0.5 and R: *F* (*x*_1_, *x*_2_) *<* −0.5. This is because *F* (*x*_1_, *x*_2_) always visits the range [−0.5, 0.5] in a perfectly synchronized manner with the interoceptive input, as shown in Fig. 2, leading to the overestimation of the expected value of the stochastic perceptual alternation shown in Fig. 2-(b). We collected the phase of the interoceptive signal *g*(*t*) or *A*(*t*) when the state of a WTA transitioned among these three states, and computed the probability density shown in Fig. 3 and 4.

#### WTA behavior driven by the periodic neuromodulator *g*(*t*)

First, we analyze how interoceptive neuromodulator *g*(*t*) affects the behavior of a WTA circuit. Note that we can assume *τ* = 1 without losing generality ^2^. If we take a Taylor expansion of Eq .1 with *x*_*i*_ ∼ 0, and if we consider a variable 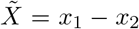, Eq. 1 is reduced to the one-dimensional linear differential equation as

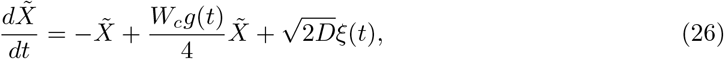

where *W*_*c*_ = *w*_*ii*_ − *w*_*ij*_ and *g*(*t*) = *g*_*b*_ + *g*_*g*_ sin (2*πft*). 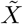 converges to 0 if *W*_*c*_*g*(*t*)*/*4 *<* 1 (conventional negative feedback mode), and diverges if *W*_*c*_*g*(*t*)*/*4 *>* 1 (positive feedback mode). Therefore, if *W*_*c*_*g*(*t*)*/*4 *>* 1, then a winner neuron can switch because the small noise-induced perturbation, represented by the variable *ξ*(*t*), is amplified by the positive feedback mode. However, if *g*_*b*_ ≫ *g*_*g*_, the winning neuron diverges ; *x*_1_ − → *x*_2_±∞ . This implies that the winning neuron’s transition is blocked.

#### WTA behavior driven by the additive input signal *A*(*t*) with normal and small activation gain *g*_*b*_

We analyze how interoceptive common input *A*(*t*) affects the behavior of a WTA circuit. If the acitivation gain *g*(*t*) in Eq. 1 is constant and small as *g*_*g*_ = 0 and *g*_*b*_∼ 0, the WTA neurons have rather linear dynamics, then Eq. 1 is reduced to the following one-dimensional dynamics with respect to 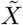 as

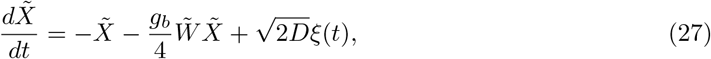

where 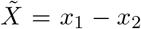 and 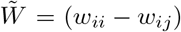. Eq. 27 indicates that in the linear WTA regime (i.e., *g*_*b*_ ≪ 1), the interoceptive common input *A*(*t*) does not contribute to the dynamics of the neural state transition. For additional supporting material, please refer to Fig. 10-(b).

Fig. 9 shows the WTA behavior driven by the additive input signal *A*(*t*) = *A*_*g*_(*t*) sin(2*πft*)+*A*_*b*_, with the normal activation gain *g*_*b*_ = 1.5 (A), and the small gain *g*_*b*_ = 0.1 (B). Note that the red line denotes *f* (*x*_1_) (the output of *x*_1_), and the blue line denotes *f* (*x*_2_). Black line in the panel (B) denotes *f* (*x*_1_) − *f* (*x*_2_), and it is shown *f* (*x*_1_) − *f* (*x*_2_) ∼ 0, and the WTA loses the bistability.

#### Detail analysis of the effect of interoceptive additive input to WTA

To analyze the effect of interoceptive input *A*(*t*) in Eq. 1, we use Itô’s formula [29]. Eq. 1 can be written in the following differential form as

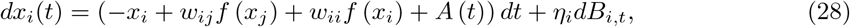

and, furthermore, in the integral form as

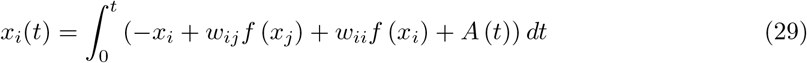

where (*i, j*) is either (1, 2) or 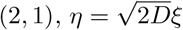 and *dB*_*i,t*_ describes the path of noise process (note that we assumed *τ* = 1 for simplicity). Because the Taylor expansion of *f* (*x*_*i*_ + *dx*_*i*_) is described as

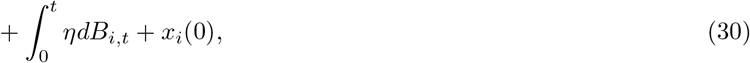

Because *df* (*x*_*i*_) = *f* (*x*_*i*_ + *dx*_*i*_) − *f* (*x*_*i*_), we have the differential form of *f* (*x*_*i*_) as follows:

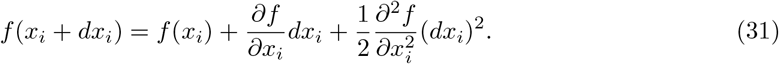

Note that in the limit *dt* → 0, (*dB*_*t*_)^2^ → 0, *dB*_*t*_*dt* → 0, *dt*^2^ → 0, and *dx*^2^ → *η*^2^*dB*_*i,t*_ [29].

By substituting Eq. 28 to Eq. 32, and taking the integral of *df* (*x*_*i*_), we have the integral form of Eq. 32

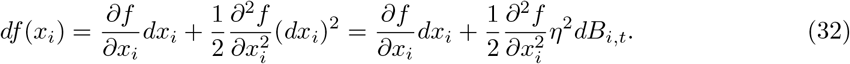

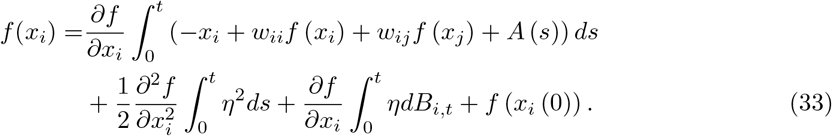

We can compute *F* (*x*_1_, *x*_2_) = *f* (*x*_1_) − *f* (*x*_2_) as

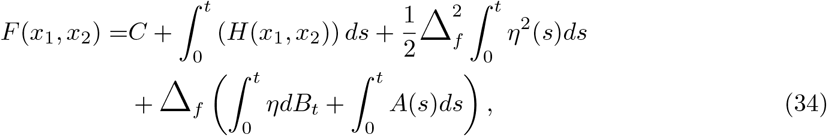

where *C* = *F* (*x*_1_(0), *x*_2_(0)), Δ_*f*_ = *∂f/∂x*_*i*_ − *∂f/∂x*_*j*_, 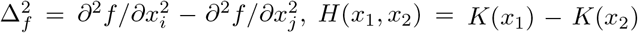, and furthermore, 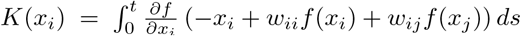*ds*. Taking the Taylor expansion of Eq. 34 around *x*_1_ = 0 and *x*_2_ = 0, we have the result shown in Eq. 3. It should be noted that the dependency of *F* (*x*_1_, *x*_2_) on *A*(*s*) disappears when we take the Taylor expansion around *g* = 0.

## C Details of simulated Iowa gambling task

In a simulated IGT (Fig. 3), we have assumed that bad deck (BD) is configured to return a large positive score but the expected return is negative *R*_*n*_ = −25 − *r*_0_ + 375*ξ*, and good deck (GD) is configured to return only a small positive score, but the expected return is positive *R*_*n*_ = 25 − *r*_0_ + 75*ξ* (*r*_0_ is bias parameter), and *ξ* is a Gaussian noise (*n* denotes the *n*-th decision). For simplicity, the IBA of an agent is described by sinusoidal activity as *g*_*g*_ sin(2*πf*_*r*_*t*) + *g*_*b*_.

The agent choses either BD or GD at every *T*_*d*_ (*T*_*d*_ is random), and receives the score *R*_*n*_. The agent internally evaluates *R*_*n*_ using the utility function as *U*_*n*_ = *U* (*R*_*n*_) = *R*_*n*_ for *R*_*n*_ ≥ 0 and 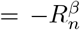 for *R*_*n*_ *<* 0 (Note that *β <* 1 denotes underestimation of risks), and the output of the utility function *U*_*n*_ is fedback to IBA parameters based on the utility function of the outcome of *n*th decision as *z*(*n* + 1) = *z*(*n*) + Δ_*z*_(*λU*_*n*_ + *η*_*n*_) + *z*(*n*)*ζ*_*n*_, where *z* denote either *f*_*r*_, *g*_*g*_, or *g*_*b*_, *η*_*n*_ and *ζ*_*n*_ is a small Gaussian noise (with standard deviation 0.05, and 0.001), and Δ_*z*_ is simply a coefficient (i.e., Δ_*z*_ is either 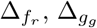, or 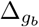). We used a scaling factor *λ* = 1*/*350. *η*_*n*_ represents additive Gaussian noise and *ζ* represents multiplicative Gaussian noise. These noise components are introduced to avoid local minimum.

The rationality in IGT is measured by 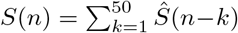, where *Ŝ*(*q*) = +1 if the decision in *q*th step is GD, and 1 − otherwise; if the agent keeps choosing GD, *S*(*n*) converges to 1, *S*(*n*) = −1 if the agent is sticked to BD, and *S*(*n*) ∼ 0 if the agent keeps wondering between the GD and BD (Fig. 3-(B), the middle panel). However, Fig. 3-(D) shows that metalearning, i.e., an appropriate setting of Δ_*z*_ is necessary for the occurrence of rationality emergence, and if Δ*z* is not adequate, the system is trapped to irrational behaviors.

### A WTA neural circuit and reward-modulated Hebbian learning

In Fig. 3, we add additional synaptic connection from a competing layer which is described by Eq. 1 to an input neuron layer. The vector of neural state **X** and the weight matrix **W** in Eq. 25 is replaced to **X** = (*x*_0_, *x*_1_, *x*_2_, *x*_3_, *x*_4_)^*T*^ (where *T* denotes transpose), and **W** is replaced to 4 × 4 matrix. The input layer neurons are assigned to *x*_0_ and *x*_1_. The weight of inhibitory connection from *x*_2_ and *x*_3_ to the input neurons *x*_0_ and *x*_1_ are fixed as *w*_20_ = *w*_21_ = *w*_30_ = *w*_31_ = −3.5. The excitatory connections from input layer neurons to neurons in a winner-take-all (WTA) layer are subject to reward-modulated Hebbian learning, and the update rule is described as

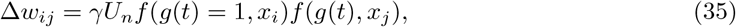

where *γ* is learning rate = 0.001 or 10^*−*5^, *U*_*n*_ is the output of utility function. For the input layer neurons, we use the fixed gain *g*(*t*) = 1. For the WTA layer neurons, we assumed that *g*(*t*) is modulated by IBA as *g*(*t*) = *g*_*g*_ sin(2*πft*) + *g*_*b*_.

## D Statistical information

We collected binocular rivalry data from 30 participants. Fig. 13 shows the number of valid participants’ data used in the TE permutation test and MCMC analysis after removing measurement errors and artifacts using the above-described methods.

**Figure 13:**
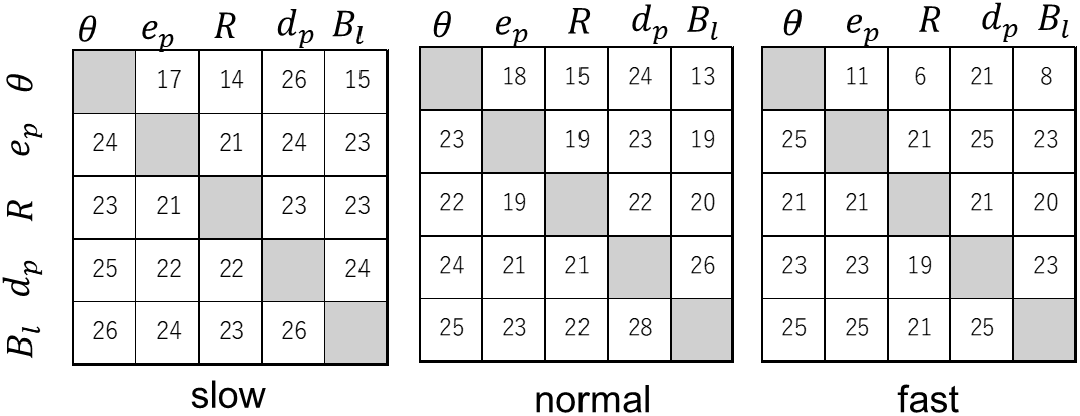
The number of valid participants’ data used in the TE permutation test

## Footnotes

1 The point process *Y* corresponding to the respiration phase *θ* with *I*_*y*_ = 10 denotes one cycle of respiration; for we computed the point process by dividing the parameter space *θ* : [0, 2*π*] into 10 subspaces.

2 The time-constant *τ* in Eq.1 can be removed by applying adequate scaling of **X** and *t*, and we have 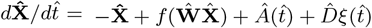, where 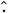*·* represent the scaled variable.

